# Coordination of local and global responses in individual mouse thalamic local interneurons

**DOI:** 10.1101/2025.10.20.683501

**Authors:** Kunyun Wang, Wenxin Hu, Yue Fei, Chen Li, Chang Liu, Ashley Ontiri, Deng Zhang, Yuanyu Chen, Michelle Luh, Liang Liang

## Abstract

Inhibitory interneurons, featuring presynaptic release sites and spiking activity in dendrites, present a puzzle in their signal organization and coordination. We investigated this in local interneurons (LINs) of the primary visual thalamus using two-photon calcium imaging in awake mice with both local and full-screen visual stimuli. Dendrites in single LINs showed consistent global responses to full-screen stimuli but varied responses to local stimuli. Dendrites from the same LIN exhibited different receptive fields that followed retinotopic progression. The similarity between dendritic and somatic responses decreased with increasing distance. Weak local stimuli could excite dendrites locally, while salient local stimuli triggered global responses. Salient global stimuli further enhanced motion selectivity, preference consistency, and peak responses. Thus, LINs leverage both local and global dendritic activation to perform spatially-organized, feature-selective, and size-dependent motion processing, providing new insights into the complexity and precision of sensory processing in the early visual system.

**Highlights:**

- Local interneurons (LINs) exhibit diverse yet sharp tuning to moving stimuli.
- display an orderly shift of receptive field centers along their dendrites.
- The propagation of calcium signals under local stimuli depends on stimulus strength.
- Full-screen stimuli enhance motion selectivity and peak responses.

## Introduction

Inhibitory interneurons are crucial for regulating sensory processing. They can shape the computation of sensory feature selectivity^1–7^, expand the dynamic range^8–10^, refine sensory receptive fields^11–13^, and enhance temporal precision and response reliability^14–17^ of their target neurons. These functions are supported by diverse types of interneurons that can display distinct morphology and employ various computational strategies.

While many interneurons exhibit classical dendrite-axon polarity, receiving inputs via dendrites and releasing outputs from axons, several types of interneurons feature pre- and post-synaptic sites distributed along their dendrites. Examples include horizontal cells and amacrine cells in the retina^18–20^, granule cells in the olfactory bulb^21^, interneurons in the superior colliculus^22^, and local interneurons (LINs) in the dorsal lateral geniculate nucleus (dLGN) of the thalamus^23–28^. The dendrites of these neurons are often thought to operate as multiplexors with many independent input-output processing units. For instance, individual dendrites of starburst amacrine cells are tuned to different centrifugal motion directions based on the branching direction of each specific dendrite^29^; proximal and distal synaptic inputs for granule cells arise from distinct sources and have different kinetics *in vitro*^30^.

Although axonless retinal interneurons typically do not fire action potentials^19,31,32^, spikes can be observed in other interneurons with dendrodendritic connections, such as granule cells and LINs^33–36^. Yet, it remains poorly understood how these neurons utilize both local input-output units and propagating spikes to process sensory signals *in vivo.* The integration of local activation can trigger spikes, and backpropagating spikes, in turn, can also influence local input-output computations. If individual dendritic compartments compute distinct visual information while global excitation pools mixed information from different dendritic compartments, dendrites would output different information under local versus global activation. Conversely, if dendrites within the same neurons have similar stimulus preferences, what would be the role of metabolically costly backpropagating spikes when local and global activation produce consistent outputs to target neurons?

dLGN LINs serve as an ideal model for studying the local and global activity of interneuron dendrites in the brain. These LINs receive most of their synaptic inputs from retinal ganglion cells (RGCs) and send outputs to thalamocortical cells (TCs) as well as other LINs^28^. Recent evidence indicates that the dLGN is not a simple relay but rather plays a more active role in sculpting the flow of visual information^37–41^. LINs have been suggested to regulate both the temporal and spatial response properties of TCs, thereby playing an important role in dLGN visual processing^10,16,35,42,43^.

Individual mouse LINs have extensive dendritic arbors that can span the entire depth and over half the width of the dLGN^28^. Hundreds of presynaptic GABA release sites (F2 terminals) are distributed along individual dendritic arbors and participate in various modes of triadic synapses with TCs and other LINs^27,28^. Individual LINs can innervate hundreds of TCs and primarily target their proximal dendrites^27,28,44^. Although LINs also provide inhibitory output through conventional axonal terminals (F1 terminals), axons account for only a small fraction of their output^28^. Their dendrodendritic synapses have been proposed to enable fast feedforward inhibition^16^ and allow a single LIN to participate in hundreds of different neuronal interactions and inhibitory processes^28^.

*In vitro,* LINs can be activated both globally and locally. In mouse dLGN slices, both synchronous stimulation of multiple retinal axons and somatic current injections trigger sodium and calcium spikes in LINs, leading to calcium elevation without decay throughout the dendrites and widespread GABA release from dendrites. This suggests the involvement of active conductance in signal backpropagation from the soma to dendrites, resulting in global activation of LINs^45^. In both rat and cat dLGN slices, LINs can release GABA when triggered by ionotropic or metabotropic glutamate receptor activation, even in the presence of TTX to block action potentials^46,47^. Furthermore, local glutamate uncaging of distal dendrites in rat LINs rarely triggers somatic firing or calcium spikes, but it can still evoke inhibitory postsynaptic currents (IPSCs) in TCs^44^, indicating that LINs can be locally activated and release GABA without global excitation. However, local calcium signals required for local GABA release have not been directly observed^48^.

*In vivo*, action potentials recorded from LINs have been used to study their visual response properties. LINs somas have been reported to exhibit contrast selectivity, motion selectivity, and push-pull receptive field structures^35,36,49,50^. However, it remains unclear whether visual responses from different dendrites within the same LIN can be compartmentalized. Additionally, how the dual-mode activation, if present, is coordinated and relates to the visual properties of LINs is still elusive. A recent study showed that the direction and orientation selectivity of LIN proximal dendrites was similar to that of the soma under full-screen stimuli^50^. However, full-screen stimuli likely triggered backpropagating spikes that synchronized dendritic activity and therefore reflected global excitation, as suggested by previous *in vitro* studies^45^.

To determine *in vivo* whether visual stimuli could locally activate LIN dendrites, how local activation may trigger global dendritic activity, and how response properties of dendritic compartments are related to each other under local stimuli and between local and global stimuli, we employed visual stimulation of different saliencies, including full-screen and local patch drifting gratings and gratings of different contrasts. We recorded visual responses in individual dendrites and somas of LINs in the dLGN shell using chronic two-photon calcium imaging in awake mice with single dendrite resolution, and we developed methods to identify dendrites belonging to the same neuron. Combining these new strategies, we elucidated the coordination of visual responses across the dendritic arbor of individual LINs and revealed the relationship between local and global responses. Our findings suggest rules by which neurons with both local dendrodendritic synapses and spiking activity may orchestrate their dendritic computation to adaptively and precisely process sensory information of varying sizes and saliency.

## Results

### Diverse and Selective Responses in LIN Somas

To measure the visual response properties of local interneurons (LINs) *in vivo*, we used a high-resolution chronic imaging technique that we recently developed for visualizing neuronal activity in the dLGN of awake, head-restrained mice (Figure 1A)^39,51,52^. We expressed the green fluorescent calcium indicator GCaMP6f in LINs by injecting Cre-dependent GCaMP6f into the dLGN of Gad2-IRES-Cre mice (Figure 1A). In these mice, Cre-recombinase is selectively expressed in LINs in the dLGN, as demonstrated by recent single-cell RNA-seq of mouse dLGN neurons^53^. We first identified the dLGN region from several neighboring thalamic regions using the stereotypic retinotopic map of visually evoked bulk epifluorescence responses to 20° Gabor-like stimuli^51^ (Figure 1B), as we also observed Gad2-positive neurons in the lateral posterior (LP) thalamus, intergeniculate leaflet (IGL), and ventral lateral geniculate nucleus (vLGN) through the implanted cranial window.

**Figure 1.**
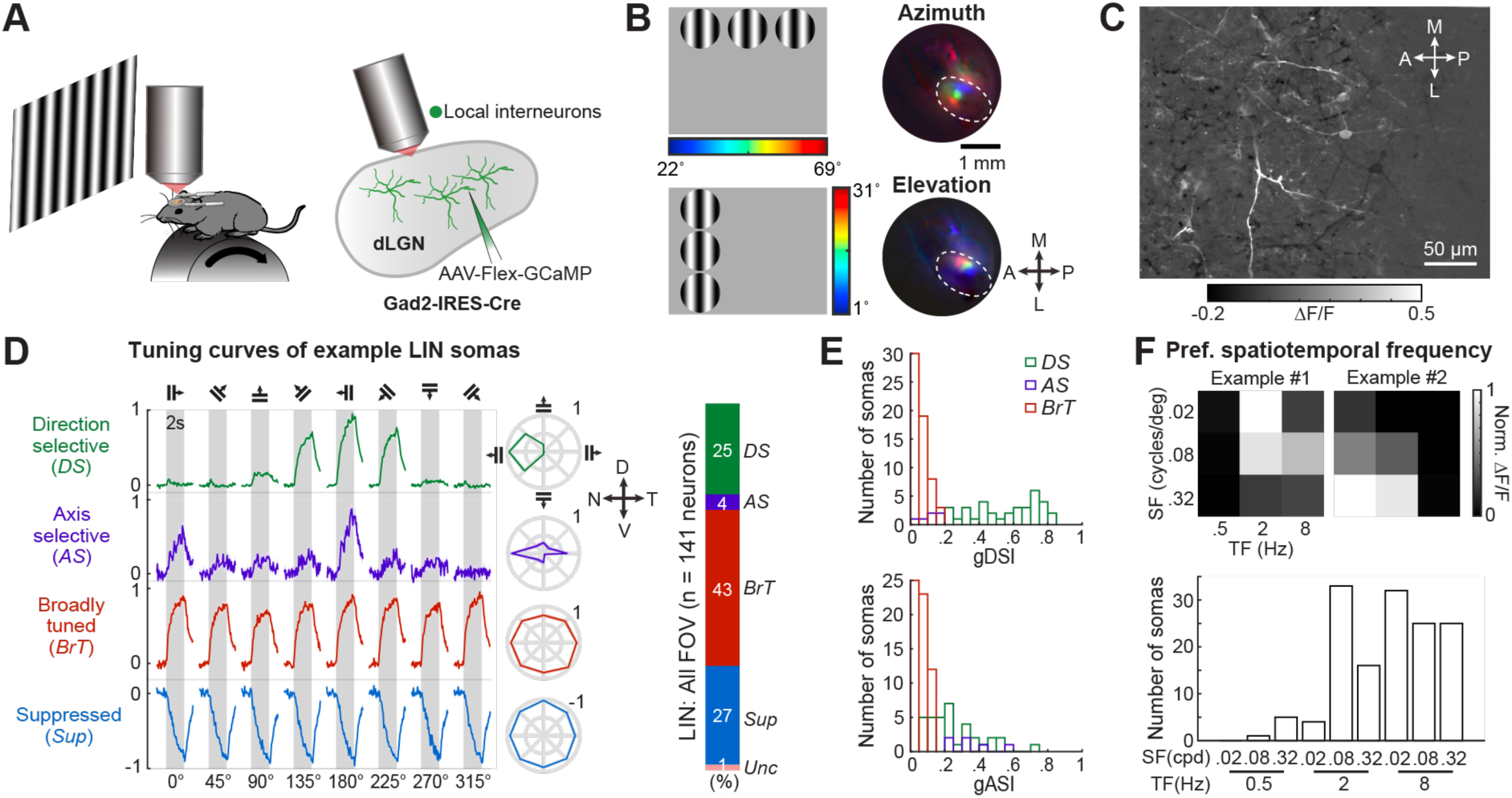
**LIN Somas Show Diverse Responses to Motion** (A) Schematic of imaging setup. *Left*, the dLGN contralateral to the stimulated eye was imaged through a cannula chronically implanted above the thalamus. *Right*, the calcium indicator GCaMP6f was specifically expressed in dLGN local interneurons in Gad2-IRES-Cre mice. (B) The dLGN region was identified by stereotypic bulk retinotopic responses from epifluorescence calcium imaging. *Left*, 20°Gabor stimuli presented at three azimuths (top) or elevations (bottom). *Right*, pseudocolored bulk calcium responses to stimuli at different azimuths (top) and elevations (bottom). White dashed line indicates the contour of the dLGN. A, anterior; P, posterior; M, medial; L, lateral. (C) Visual responses of LINs in an example field of view (FOV), presented as the sum of maximum and minimum responses across multiple stimulus conditions. ΔF/F: fractional change in fluorescence. (D) *Left*, example response timecourses (mean ± SEM) to full-screen drifting gratings for one LIN soma in each functional category (gray bars, 2s presentation of drifting gratings). *Middle*, normalized mean response tuning curves. *Right*, the proportion of somas in each category across 11 FOV from 10 mice (N = 141 somas; direction-selective [*DS*]: 24.8%; axis-selective [*AS*]: 4.3%; broadly-tuned [*BrT*]: 42.6%; suppressed [*Sup*]: 27.0%). Rarely, somas were responsive but not classified (Unclassified [*Unc*]:1.4%). (E) Distribution of the global direction selectivity index (gDSI) and the global axis selectivity index (gASI) of *DS*, *AS* and *BrT* somas. (F) *Top*, response heatmaps of two example LIN somas preferring different spatial and temporal frequency combinations. *Bottom*, distribution of preferred spatial and temporal frequency combinations of all significantly responsive LIN somas (N = 141 somas from 11 FOVs, 10 mice).

We then measured visual responses of LINs at cellular resolution using two-photon calcium imaging in the dorsal dLGN, likely the shell region^54,55^. To examine the motion selectivity and receptive field properties of LINs, we presented a battery of visual stimuli, including full-screen drifting gratings of 8 directions (45° apart), 3 spatial frequencies, and 3 temporal frequencies, and 5° x 40° bars containing spatiotemporal noises. In each field of view (FOV), we observed sparse labeling of LINs and heterogeneous responses across LINs (Figure 1C).

We first characterized somatic responses, which reflected global calcium signals. Based on responses to full-screen drifting gratings, we classified LINs into four functional categories: direction-selective (*DS*; preferring one direction of motion), axis-selective (*AS*; preferring opposite directions of motion along the same axis, also known as ‘orientation-selective’)^51,56^, broadly-tuned (*BrT*; broadly responsive across all directions), and suppressed (*Sup*; suppressed during visual stimulation)^57–59^ (Figure 1D). This classification is similar to how we categorized retinal ganglion cell (RGC) axonal boutons in the dLGN^51^ and consistent with a recent study of LINs^50^. Interestingly, the *DS* and *Sup* LINs each comprise significant fractions of the LIN population (24.8 ± 2.8% and 27.0 ± 4.5%, respectively), and a large fraction of *DS* LINs exhibited high direction selectivity (Figure 1E). In addition to displaying selective preferences for motion directions and/or axes, individual LINs also exhibited diverse yet sharp preferences for the spatial and temporal frequencies of drifting gratings (Figure 1F), a response property not examined in previous studies^36,50^. These observations reveal that LIN somas can exhibit surprisingly sharp tuning to motion information and suggest that at least a sub-population of LINs may selectively sample inputs from retinal axons.

### Comparison of Response Selectivity Between LINs and RGCs

Given that LIN somas exhibited diverse functional categories similar to RGC boutons^51^, we sought to examine whether afferent retinal visual information is biasedly sampled by LINs at the population level. To address this, we compared the response properties of LINs and RGCs input by recording visually evoked calcium responses from GCaMP6f-expressing retinal axonal boutons projecting to similar depths of the dLGN using the same set of visual stimuli (Figure S1A). Consistent with our previous findings, RGC boutons could be well-characterized by the four categories^51^. We observed that the fractions of *DS*, *AS*, *BrT* and *Sup* LINs (*DS*, 24.8 ± 2.8%; *AS*, 4.3 ± 2.1%; *BrT*, 42.6 ± 4.8%; *Sup*, 27.0 ± 4.5%) were close to that of the RGC axons projecting to the dLGN shell (*DS*, 18.7 ± 1.4%; *AS*, 10.8 ± 1.5%; *BrT*, 30.4 ± 1.7%; *Sup*, 23.7 ± 1.8%). Moreover, RGC boutons consistently preferred higher temporal frequencies, similar to LIN somas, but showed a clear preference for higher spatial frequencies.

### Identifying Dendrites from the Same LIN

When stimulating LINs with full-screen drifting gratings, we observed highly synchronized calcium activity throughout the dendrites of the same LINs. This activity synchronization likely results from action potentials that are generated at the axon initial segment and backpropagate to the dendrites via active dendritic conductance, as suggested by electrophysiological recording and imaging of LINs in acute dLGN slices^45^. To assess whether distinct dendritic domains of the same LIN exhibit functionally related responses and how these dendritic responses relate to somatic responses, especially in LINs with high motion direction selectivity, we presented local 10°- or 20°-diameter patches of drifting gratings (Gabor-like patch gratings; see Methods) at different monitor locations. Notably, distinct dendrites from the same neuron showed their peak responses to different local patch gratings, suggesting functional heterogeneity among dendrites within a single LIN (Figure 2A, top row).

**Figure 2.**
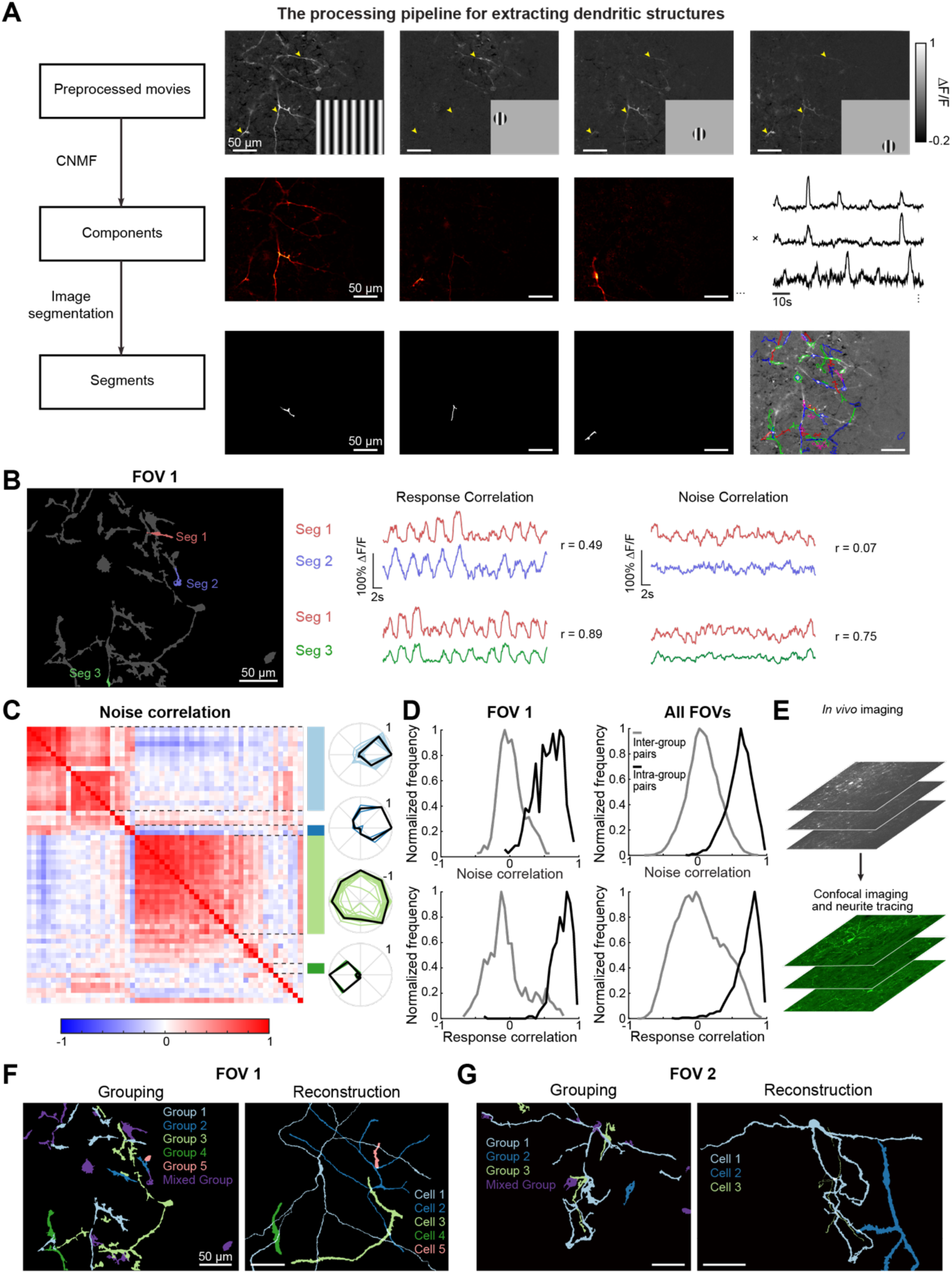
**LIN Dendrites Are Automatically Extracted and Grouped into Putative Neurons** (A) The processing pipeline for extracting dendritic structures includes (i) running a constrained nonnegative matrix factorization (CNMF) algorithm on concatenated movies under both full-screen and local patch visual stimuli to obtain spatial and temporal components and (ii) segmenting spatial components based on their spatial overlaps and connectivity. Arrowhead, dendrites from the same LIN. (B) LIN dendritic segments with high noise correlation (a measure of the pairwise similarity in the trial-to-trial fluctuation of visual responses) were grouped into the same neuron (e.g., segments 1 and 3). Dendritic segments with high response correlation but low noise correlation were separated into distinct LINs (e.g., segments 1 and 2). (C) *Left*, matrix of pairwise noise correlation for segments from FOV 1, sorted using hierarchical clustering. Four distinct blocks of segments assigned to four different neurons are highlighted. *Right*, peak-normalized tuning curves for all segments from the four neurons. (D) Histogram of the noise (top) and response (bottom) correlation for pairs of dendritic segments grouped to the same LIN (black) or different LINs (gray) from an example FOV (left) and all FOVs (right; 10 FOVs from 8 mice). (E) Schematic illustrating *post hoc* processing. After *in vivo* two-photon imaging, the mouse brain was sectioned parallel to the imaging plane. LIN neurites were then traced from the confocal stacks of the brain section containing the imaged neurons. (F-G) Results from noise-correlation-based grouping of dendritic segments (left) and *post hoc* reconstruction of individual LINs (right) in example FOVs 1 (F) and 2 (G). The same neurons are labeled with the same color.

To systematically identify dendritic segments from the same neuron and determine the extent of heterogeneity among them, we developed an algorithm to automatically extract and classify dendritic segments into distinct LINs. By combining constrained non-negative matrix factorization (CNMF) and image spatial segmentation, we extracted dendritic segments based on their distinct spatial and temporal features (Figure 2A, see Methods)^60–62^. The segments were then clustered into different neurons based on their pairwise noise correlation under full-screen drifting grating stimuli, as single-trial calcium signals are highly consistent across dendrites of the same neuron under full-screen stimuli^50^. Noise correlation calculates the correlation of trial-to-trial fluctuations around the mean response to the same visual stimulation between dendritic segments. Segment pairs from different neurons could exhibit high correlation in visual responses but low noise correlation (Figure 2B and Figure S2A), whereas pairs from the same neuron were highly correlated in both visual responses and noise (Figure 2B and 2D). This method effectively separated segments from different neurons that exhibited similar motion preferences under full-screen stimuli.

To confirm the dendritic grouping results from the algorithm, we first analyzed the visual responses of dendritic segments assigned to the same neuron and observed highly consistent visual tuning properties within each group (Figure 2C and 2D). We also performed *post hoc* neurite tracing and reconstruction of individual LINs from confocal stacks of the horizontal brain section that contained the LINs recorded from *in vivo* imaging (Figure 2E). The results of dendrite identity showed high consistency between the dendritic grouping based on *in vivo* imaging and the *post hoc* neuron reconstruction (Figures 2F, 2G, and S2B-S2D).

### Orderly Shifts in Receptive Field Centers Along Dendritic Arbors of the Same LIN

To identify the pattern of heterogeneous responses among different dendritic segments of the same LIN, we compared their responses across various visual stimuli. As expected, dendrites of the same LINs exhibited highly consistent direction preferences under full-screen drifting gratings, as illustrated in an example LIN (Figure 3B). Responses to local drifting gratings from different dendritic regions also showed similar direction preferences (strong responses to nasal-to-temporal motion and no significant responses to motion in the opposite direction) but exhibited distinct receptive fields (Figure 3C).

**Figure 3.**
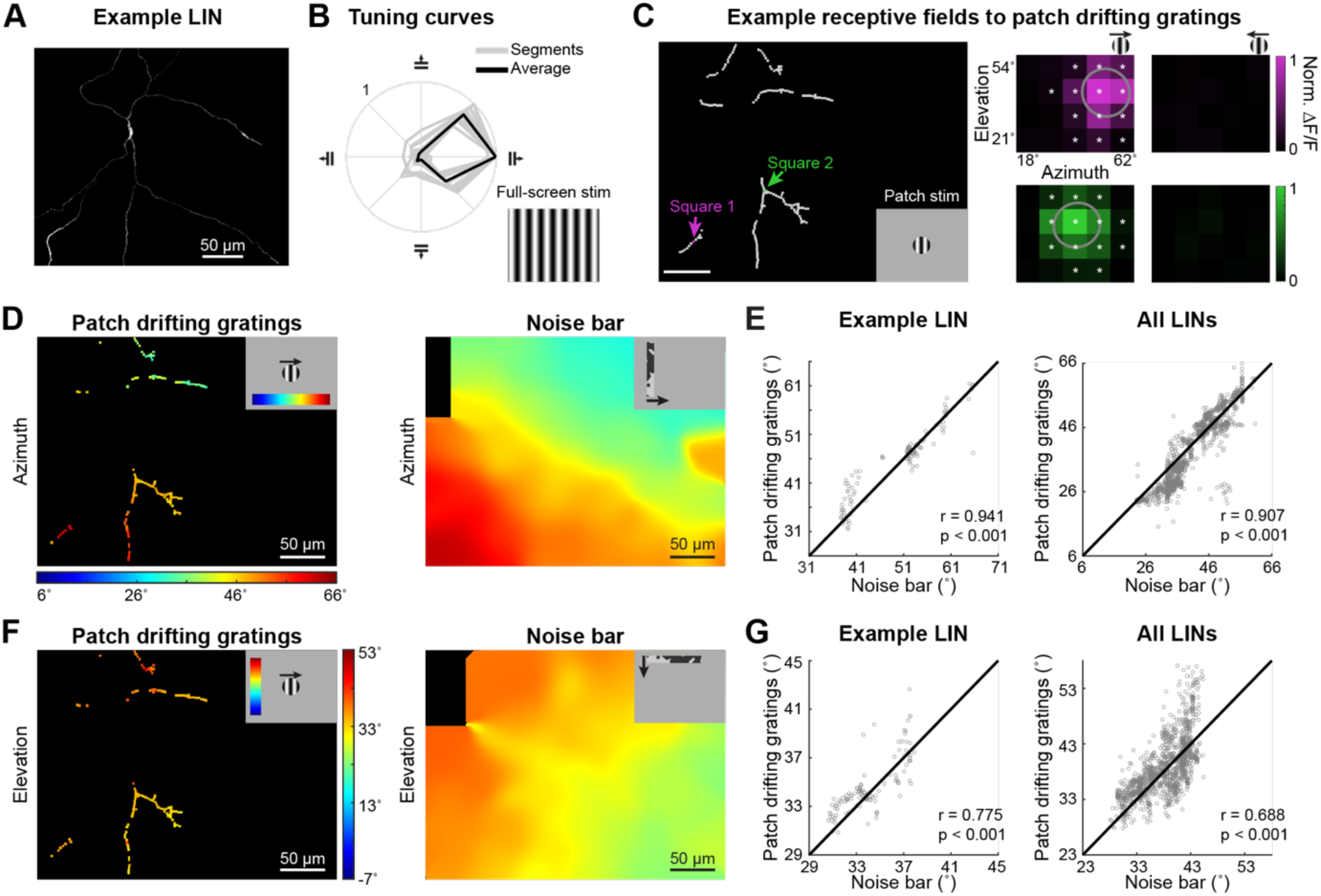
**Receptive Fields of Dendrites within a LIN Follow Retinotopic Progression** (A) Maximal Z projection of an example LIN traced and 3D reconstructed from a *post hoc* confocal image stack. (B) Different dendritic segments in the example neuron in A showed similar normalized direction-selective tuning curves to full-screen drifting gratings. Gray, tuning curves of individual segments; black, average. (C) Two example dendritic square regions of interest (ROI) from the same example LIN (shown in A and B) exhibited different receptive fields to patch drifting gratings when stimulated at their preferred direction of full-screen drifting gratings. Additionally, these ROIs showed no responses to patch drifting gratings at their null direction of full-screen drifting gratings. Gray outlines the fitted receptive fields by 2D Gaussian functions. Asterisks indicate that the ROI exhibited significant and reliable responses to the patch drifting gratings presented at the specific location and direction. (D) *Left*, azimuth positions of receptive field centers of individual dendritic square ROIs in the example LIN. *Right*, the LIN population retinotopic progression of the FOV along the azimuth axis, mapped by vertical noise bars displayed at different azimuths and calculated after removing pixels of the example neuron (see Methods). (E) Azimuth receptive field centers of dendritic square ROIs mapped by patch drifting gratings showed a high correlation with the retinotopy mapped by vertical noise bars in the example LIN (left) and all LINs (right, 10 neurons from 9 FOVs, 8 mice; the example LIN azimuth r = 0.941, p < 10^-53^; all LINs azimuth: r = 0.907, p < 10^-307^). (F) *Left*, elevation positions of receptive field centers of individual dendritic square ROIs in the example LIN. *Right*, the retinotopic progression of the FOV along the elevation axis, mapped by horizontal noise bars and calculated after removing the pixels of the example neuron. (G) Elevation receptive field centers of dendritic square ROIs mapped by patch drifting gratings showed a high correlation with the retinotopy mapped by horizontal noise bars in the example LIN (left) and all LINs (right, 10 neurons from 9 FOVs, 8 mice; the example LIN elevation: r = 0.775, p < 10^-23^; all LINs elevation: r = 0.689, p < 10^-195^).

To determine the relationship between the receptive fields of different dendritic regions, we divided the dendrites of the same LIN into equal-sized square regions of interest (ROI) and mapped the progression of receptive fields along the dendritic arbor at a high spatial resolution. For each dendritic square ROI, we measured its receptive field using local grating stimuli at the neuron’s preferred direction (e.g., the 0° direction for the example neuron shown in Figure 3D). We found that the preferred receptive field locations were significantly different across dendritic square ROIs from the same LIN but were arranged in an orderly manner along both the azimuth and elevation axes (Figure 3D and 3F left).

Furthermore, we calculated the population retinotopic map of the FOV along the azimuth and elevation axes by mapping the preferred locations of the vertical or horizontal noise bars for local population averages (including both LIN neurons and neuropils but not the single LIN of interest) (Figure 3D and 3F right). The vertical and horizontal noise bars both contained a wide range of spatiotemporal signals and were effective in eliciting responses from many LINs. We observed a coarse-scale retinotopic organization over the LIN population, similar to the populational retinotopic map of RGC axons^51^. Importantly, we found that the receptive field centers of dendritic domains from single LINs were highly consistent with the population retinotopic preferences for the example LIN and for all single LINs pooled from 10 FOVs of 8 mice along both azimuth and elevation axes (Figure 3E and 3G).

These data indicate that different dendrite regions within a LIN receive distinct retinal inputs preferring different parts of the visual field, and these inputs are organized in a precise manner consistent with the population retinotopic progression of LINs. Dendrites from a single LIN can span nearly 400 μm in breadth along the medial-lateral axis of the dLGN^28^, corresponding to ∼40° along the azimuth axis in the visual space of RGC boutons^51^. Our results showed that the dendritic respective fields within a single LIN are not disordered but maintain high consistency with their local neighbors across the large span of the dendritic arbor. As LIN dendrites can directly release GABA in the absence of action potentials via F2 terminals when tested in brain slices^44^, the retinotopic organization of dendritic responses suggests that dendrites of a LIN can provide retinotopically consistent visual information to their local target neurons if these dendrites are locally activated *in vivo*.

### Propagation of Calcium Signals Under Local Stimuli

We next compared the receptive field sizes of the dendrites, soma, and axon from the same LIN under an identical set of local drifting gratings, as the somatic and axonal activity to local stimuli may reflect the global responses of LINs to these stimuli. Local stimuli could evoke somatic calcium responses (Figure 4B). Moreover, we found that somas responded to more positions than did individual dendritic domains (Figure 4C; additional results for a second example LIN are shown in Figure S3B and S3C), and somatic receptive fields resembled the summed dendritic receptive fields (Figures 4B and S3B), indicating that somas integrated dendritic responses to local stimuli. However, somas also filtered out weak dendritic responses, as stimulation presented on the periphery of the summed dendritic receptive fields failed to drive significant somatic responses (Figures 4B and S3B).

**Figure 4.**
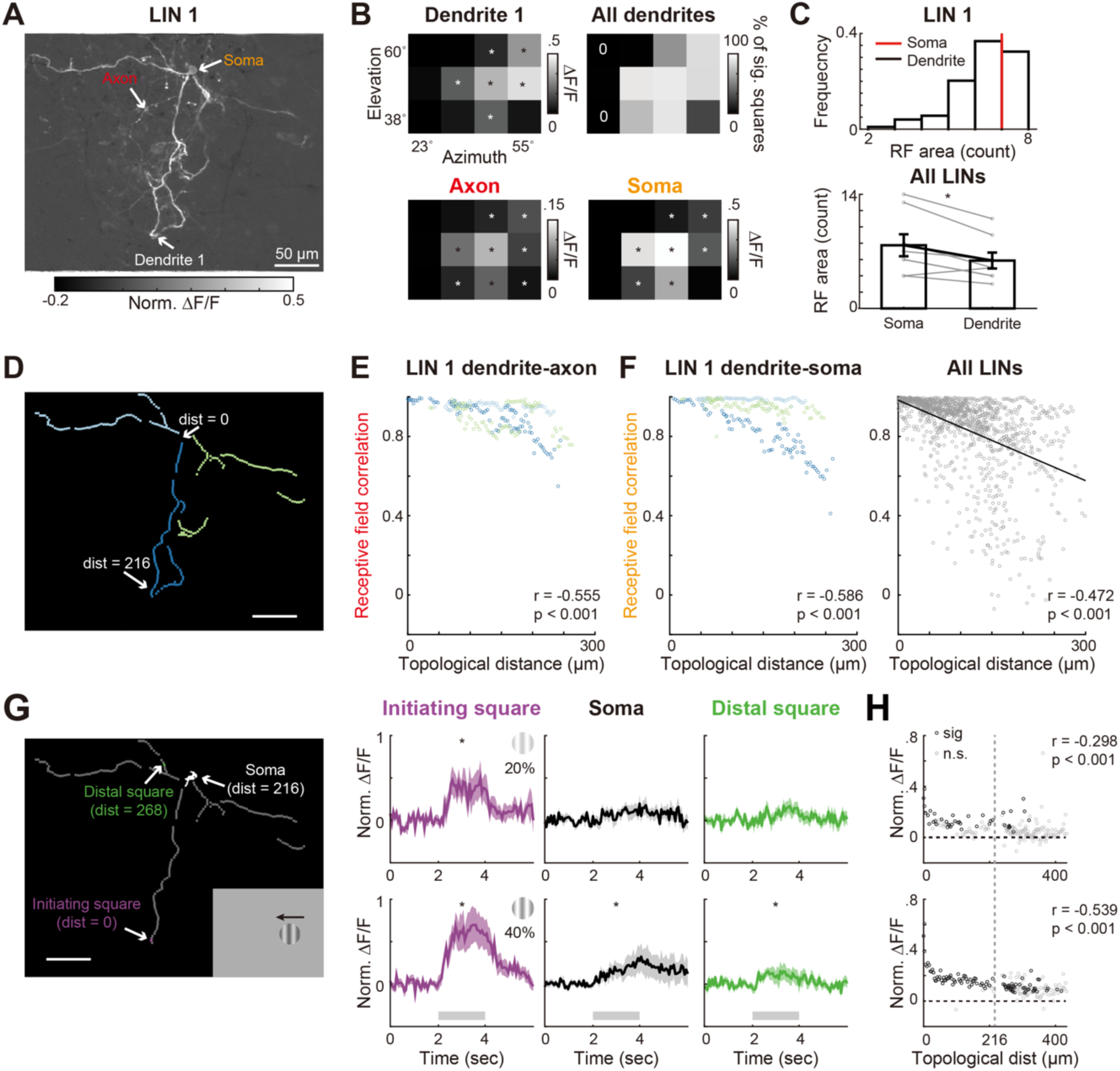
**Comparison of Receptive Fields of the Dendrites, Soma, and Axon from the Same LIN.** (A) The sum of maximum and minimum visual responses across multiple stimulus conditions in an example LIN (LIN 1). An axon-like structure (with a thinner shaft than the dendrites and *en passant* bouton-like structures) was grouped together with dendrites of LIN 1. (B) Visual responses to patch drifting gratings presented at 4 azimuth x 3 elevation locations from an example dendritic square ROI (*Top-left*), the axon (*Bottom-left*), and the soma (*Bottom-right*) of LIN 1. Asterisks (*) indicate the stimulus positions where patch stimuli evoked significantly positive and reliable responses. *Top-right*, the percentage of dendritic square ROIs from LIN 1 that exhibited significant visual responses to patch stimuli at a given location, with ‘0’ indicating no significant ROIs responsive to the location. (C) *Top*, histogram of receptive field (RF) areas (defined as the number of locations where an ROI exhibited significant and reliable responses to patch stimuli) for the dendritic square ROIs and soma of LIN 1. *Bottom*, receptive field areas of somas and medians of receptive field areas of dendritic square ROIs across all LINs with a large fraction of their dendrites observed in the imaging FOV. Somas had larger receptive fields than the medians of dendrites for these LINs (Wilcoxon signed-rank test, p = 0.031, N = 8 LINs). (D) Three major dendritic branches (each a separate color) of LIN 1. (E) Dendrite-axon receptive field correlations as a function of the topological distances of dendritic square ROIs from the axon. Receptive field correlations were negatively correlated with the topological distances of dendritic square ROIs from the axon (Pearson correlation: r = −0.555, p < 10^-19^; N = 231 dendritic square ROIs). Results are color-coded by the dendritic branch identity as labeled in D. (F) Dendrite-soma receptive field correlations as a function of the topological distances of dendritic square ROIs from the soma of LIN 1 (*left*) or somas across all LINs (*right*). Black, linear regression of the scatter plot. Receptive field correlations were negatively correlated with the topological distances of dendritic square ROIs from the soma (Pearson correlation: LIN 1, r = −0.585 p < 10^-21^, N = 231 dendritic square ROIs; All LINs, r = −0.468 p < 10^-64^, N = 1,171 dendritic square ROIs from 8 FOVs in 7 mice). Note that only dendritic square ROIs displaying significant and reliable responses to at least one condition of patch drifting gratings were included for E and F. (G) *Left*, masks of LIN 1 with the initiating dendritic square ROI, a distal dendritic square ROI from another major branch, and the soma highlighted. *Right*, average normalized response traces of the ROIs to patch drifting gratings of 20% (*top row*) and 40% (*bottom row*) contrasts (mean ± SEM). Asterisks (*) indicate the responses are significantly positive and reliable. While all three ROIs showed significant responses to 40% contrast patch drifting gratings, only the initiating square responded significantly to 20% contrast, suggesting that calcium signals propagated across the soma under 40% contrast but not 20% contrast. (H) Scatter plot of normalized responses from ROIs of the LIN against these ROIs’ topological distances from the initiating square under 20% contrast (*top*) and 40% contrast (*bottom*) patch stimuli. Normalized responses and topological distances were negatively correlated under both contrasts (20%: r = −0.298, p < 10^-5^; 40% contrasts: r = −0.539, p < 10^-17^), suggesting calcium signals propagated with decay.

Furthermore, we calculated the spatial correlation of receptive fields between the soma and dendritic domains to assess their receptive field similarity. This correlation decreased with increasing topological distance of the dendrites from the soma (Figure 4D and 4F). Interestingly, different dendritic branches of the same LIN exhibited distinct rates of decrease (Figure 4F, branches are denoted by different colors), suggesting distinct intrinsic biophysical properties, such as membrane properties and ion channel distributions, for signal conduction across different dendritic branches. Additionally, we captured an axon-like structure with a thinner shaft and *en passant* boutons in one LIN (Figure 4A). The receptive field correlation between dendritic domains and the axon initial segment similarly decreased with increased topological distance (Figure 4E).

Since local stimuli at the peripheral receptive field could significantly evoke responses in dendritic domains but not in somas (Figures 4B and S3B), we conjectured that manipulating stimulus strengths would locally activate dendrites or globally activate the LIN. Therefore, we tested how LIN dendrites responded to local stimuli with different contrasts. We predicted that low-contrast local stimuli could evoke local dendritic calcium signals, whereas high-contrast local stimuli could evoke strong local dendritic signals that trigger somatic firing and result in global dendritic calcium signals. Our observations supported this prediction. We presented local drifting gratings under 20% and 40% contrasts at different locations tiling the LIN receptive fields. For each stimulation location, we identified an initiating dendritic square ROI if this ROI consistently showed the strongest response under different contrasts (see Methods). We found that when some distal dendrites served as the initiating dendrites for local visual stimuli, responses to low-contrast local stimuli significantly attenuated before reaching the soma, whereas responses evoked by higher-contrast local stimuli could evoke somatic responses (Figure 4G; see also Figure S3D-S3E for results of another LIN). Moreover, there was a significant decrease in the normalized response magnitude with increased distance from the initiating segments (Figures 4G and S3F), indicating a decay of propagating calcium responses from the initiating ROI to the soma.

In conclusion, LIN somas integrate and threshold dendritic responses to local stimuli, depending on the strength of dendritic responses and topological distances between dendritic domains and the soma. While local weak stimuli can engage LIN local dendrites to affect local microcircuits, local salient stimuli can evoke global responses of the LIN to influence many more neurons in the dLGN through GABA release from its large dendritic arbor.

### Enhanced Direction Selectivity and Preference Consistency in LIN Dendrites by Full-Screen Stimuli

In the example *DS* LIN, we observed consistent preferred directions between full-screen and local stimuli when recording responses to two opposite motion directions (Figure 3B and 3C). Here, we tested whether the preferred direction and direction selectivity remained consistent between full-screen and local stimuli when multiple motion directions were presented to evoke responses in LINs. For each dendrite, the preferred directions under local stimuli were derived from its optimal patch stimulation location (see Methods).

As shown in an example dendritic segment (Figure 5A and 5B), the preferred motion directions under full-screen and local drifting gratings were the same, but full-screen stimuli led to a stronger peak response and a sharper direction tuning curve. Across dendritic segments within the same example neuron, the median preferred motion directions were not significantly different under full-screen and local stimuli. However, local stimuli resulted in more dispersed preferred directions and larger tuning widths compared to full-screen stimuli (Figure 5C). Furthermore, under full-screen stimuli, response magnitudes at the preferred directions were significantly enhanced, while responses at orthogonal and null directions were suppressed, across dendritic segments of the example neuron (Figure 5D).

**Figure 5.**
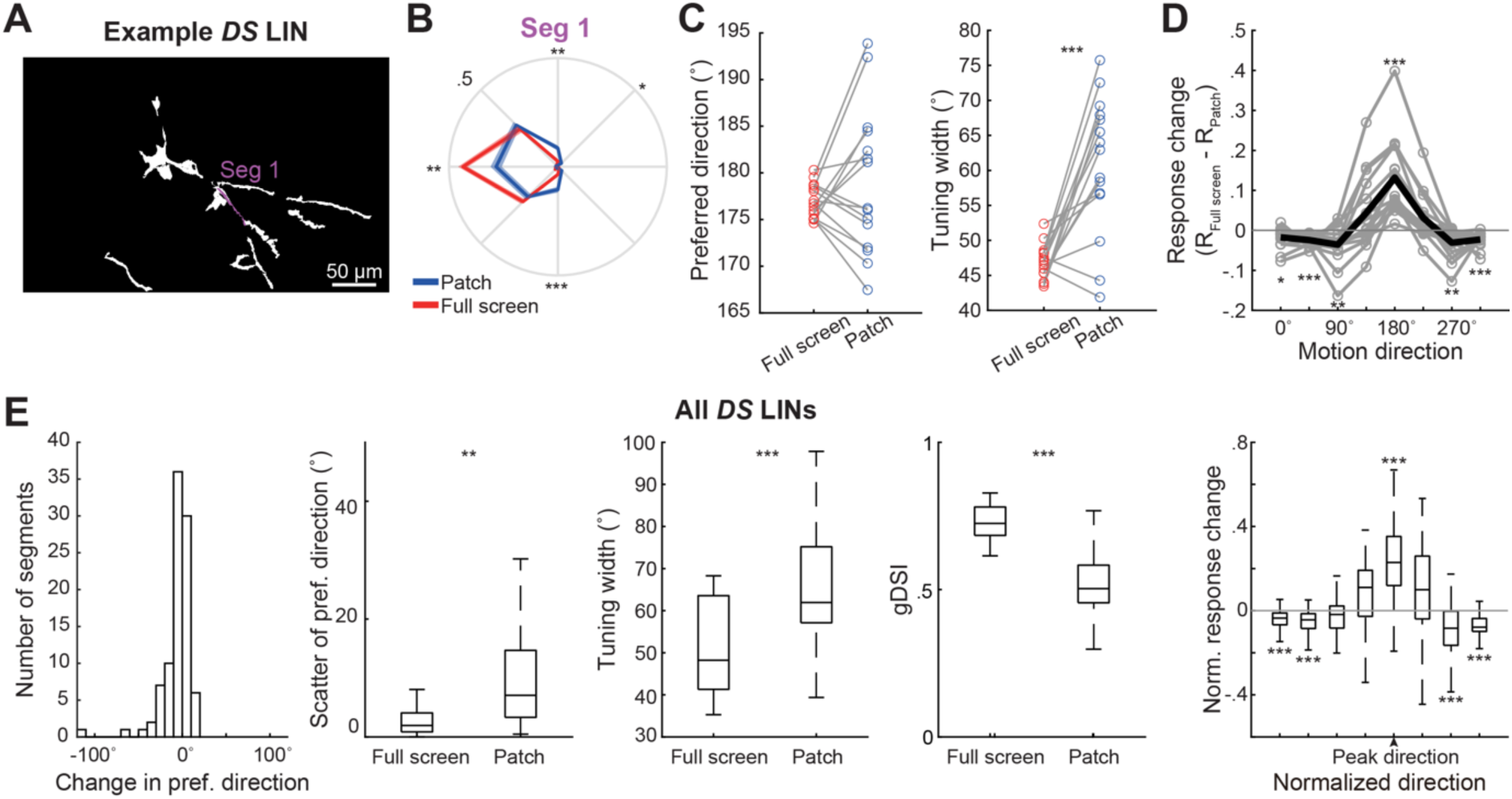
**Full-Screen Stimulation Enhances Tuning Selectivity in *DS* LIN Dendrites.** (A) Dendritic segments from an example direction-selective (*DS*) LIN with a segment (Seg 1) highlighted. (B) Tuning curves of Seg 1 to full-screen and patch drifting gratings, both at 80% contrast. (C) *Left*, the median preferred directions of all dendritic segments to 80%-contrast full-screen and patch drifting gratings from the example neuron were not significantly different (Wilcoxon signed-rank test, p = 0.421). *Right*, tuning widths of all segments from the example neuron to full-screen drifting gratings were significantly smaller than those to patch drifting gratings (Wilcoxon signed-rank test, p < 10^-3^). N = 15 segments. (D) Differences in responses (ΔF/F) to 80%-contrast full-screen and patch drifting gratings at each direction. Results from individual segments are connected by gray lines. *Black*, the average across all segments. Responses to 180° (preferred direction of the neuron) were significantly enhanced (p < 10^-3^). Responses to 0° (p = 0.013), 45° (p < 10^-3^), 90° (p = 0.004), 270° (p = 0.004), and 315° (p < 10^-3^) directions were significantly suppressed. Responses to 135° (p = 0.229) and 225° (p = 0.064) were not significantly changed. (E) Comparison of dendritic response properties under full-screen versus patch drifting grating stimuli for all *DS* LINs with a large fraction of their dendrites observed in the imaging FOV. The preferred directions between the two types of stimuli (𝐷𝑖𝑟*_Full screen_* – 𝐷𝑖𝑟*_Patch_*) showed small differences across dendritic segments (p = 0.073). Preferred directions exhibited significantly less variation under full-screen stimuli (p = 0.002). Full-screen drifting gratings also resulted in significantly smaller tuning widths (p < 10^-8^) and significantly higher gDSI (p < 10^-24^). Normalized response changes between full-screen and patch drifting gratings, calculated as (𝑅*_Full screen_* − 𝑅*_Patch_*)/(max (|𝑅*_Full screen_* |) + max (|𝑅*_Patch_*|)), showed significantly enhanced responses at the preferred direction (measured under full-screen drifting gratings) but significantly suppressed responses to null directions (peak direction, p < 10^-18^; −45°, p = 0.067; −90°, p = 0.054; −135°, p < 10^-14^; −180°, p < 10^-11^; +45°, p = 0.201; +90°, p < 10^-3^; +135°, p < 10^-21^). N = 6 neurons from 6 FOV, 6 mice. A linear mixed-effects model was used for comparisons of pooled results. *, p < 0.05; **, p < 0.01; ***, p < 0.001.

Consistent findings were observed across *DS* LINs. Although the average preferred directions of dendritic segments did not significantly differ between full-screen and local stimuli (Figures 5E and S4A), the preferred directions of dendrites under full-screen stimuli were less scattered than under local stimuli (Figures 5E and S4B). Moreover, dendritic tuning widths were smaller, and gDSIs were higher under full-screen stimuli compared to local stimuli (Figures 5E, S4C, and S4D). Sharpened direction tuning with increased stimulus size was also observed among direction-selective LIN somas and RGC axonal boutons in the dLGN (Figure S5), suggesting that RGC inputs contribute to the stimulation size-dependent direction selectivity of LINs.

We then compared the response magnitudes at each motion direction between full-screen and local stimuli. With increased stimulus size, LIN dendritic responses to their preferred directions were significantly enhanced in 5 out of 6 neurons, while their responses to the null (opposite) directions were significantly suppressed in all 6 LINs (Figure S4E). The increase in visual stimulation size led to significant decreases in responses at the null and orthogonal directions in *DS* LIN somas and *DS* RGC boutons (Figure S5 B_3_ and D_3_). Notably, *DS* RGC boutons were also significantly suppressed at their preferred and neighboring directions (Figure S5 D_3_). These results suggest that the selective enhancement of LIN dendritic responses at the preferred directions under full-screen stimuli is not directly inherited from individual RGC inputs. Instead, it likely results from the summation of synchronized inputs from all dendrites of the neuron and the triggering of strong global calcium activity and active dendritic mechanisms, which overrides the surround suppression of individual RGC inputs. Such enhanced responses at the preferred direction also contributed to the increased direction selectivity of *DS* LINs.

In summary, with increased stimulus size, the direction selectivity of *DS* LIN dendrites was enhanced by increased responses at the preferred direction and decreased responses at the null direction. Additionally, preferred directions were more consistent across dendritic segments under full-screen stimuli, suggesting that dendritic responses to global stimuli are more mutually influential, enabling stronger coordination among different dendrites, possibly through backpropagating action potentials.

### Factorization of LIN Dendritic Responses to Local Stimuli

Having established that dendrites within a LIN showed compartmentalized responses to local stimuli, we asked whether dendritic responses could be explained by multiple response components differentially weighted by distinct parts of the dendritic arbor. We decomposed the dendritic responses to local stimuli into the product between response components and dendritic domain weights, using non-negative matrix factorization (NNMF)^63^ (Figure 6A). Real responses of each dendritic square ROI could be reconstructed as a weighted sum of these response components. To determine the optimal number of components, we used a cross-validation procedure for matrix factorization (see Methods)^63^, where we repeatedly performed the factorization on different training sets 200 times for an increasing number of response components (Figure 6B) and calculated the mean squared errors (MSEs) of the model on testing sets. To avoid losing generalizable information and overfitting, we chose the model with 5% more error than the minimal error.

**Figure 6.**
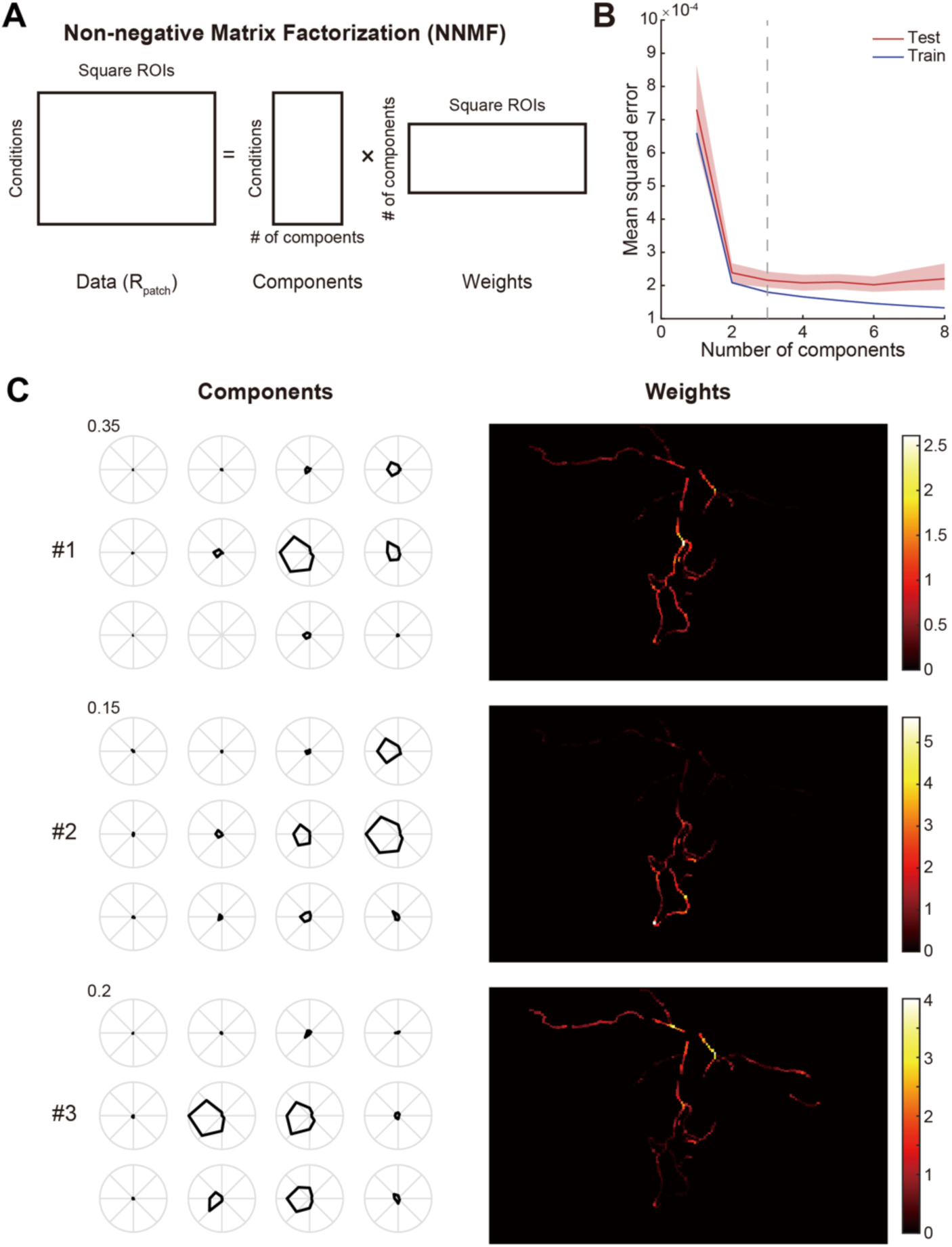
**Factorization of LIN Dendritic Responses to Local Stimuli.** (A) Schematic of non-negative matrix factorization of LIN dendritic responses. (B) Mean squared error (MSE) versus the number of response components for training (blue) and test (red) datasets in cross-validation (N_repeats_ = 200; lines, mean; shaded areas, the 95% confidence intervals; dashed line, the elbow point determined by the location of the test MSE closest to 5% above its minimal value). (C) *Left*, response patterns (responses to different directions and stimulus locations) of each component. *Right*, weight distribution across dendritic square ROIs for each response component.

Three response components were obtained from the decomposition for the example LIN (Figure 6B and 6C). The three components exhibited distinct response patterns, including preferred stimulation locations and the associated tuning curves. The corresponding weight distribution showed different spatial patterns along the dendritic arbor, with continuous, smooth, but distinct dendritic domains for each response component. Multiple response components were identified for each of the other LINs (Figure S6). Matrix factorization confirmed that dendritic responses to local stimuli could be explained by multiple latent components, consistent with our observation that dendrites within one LIN could have different receptive fields and slightly different tuning curves.

## Discussion

Using high-resolution imaging to resolve calcium signals from individual dendrites in the thalamus and a range of global and local stimuli, we observed both globally and locally evoked dendritic activity within individual LINs. Under full-screen stimulation, LINs exhibited diverse responses and sharp selectivity to all tested visual features, and their dendrites showed highly synchronized responses. Under local stimulation, dendrites from the same LIN displayed different receptive fields consistent with the overall retinotopic progression in the FOV, suggesting that a single LIN pools distinct local inputs tuned to different spatial locations. The preferred directions of dendrites under local and full-screen stimuli were largely similar, indicating coordinated direction-selective inputs within single LINs. However, full-screen stimuli elicited dendritic responses with more consistent preferred directions, enhanced direction selectivity, and increased peak amplitudes compared to local stimuli. Non-negative matrix factorization confirmed the presence of multiple latent components in dendritic responses to local stimuli. Furthermore, low-contrast local visual stimuli could evoke local dendritic activity and graded calcium signals, while more salient stimuli triggered global responses. These findings suggest that LINs selectively arrange inputs across extensive dendritic arbors, utilize their unique dendritic organization to participate in both local and global visual computation, and provide inhibition with distinct spatial patterns and varying strengths according to the saliency and size of the visual stimuli.

### Potential Convergence Rules of Retinal Inputs onto LINs

#### Retinal Origin of Diverse Responses of LIN Somas to Motion in the dLGN

Our population imaging results of LIN somas and RGC boutons show that LIN somas can be well-categorized into four functional classes, similar to RGC boutons (Figures 1 and S1), suggesting that LINs potentially inherit their diverse response properties from the retina. The observation that LINs exhibited sharp and selective preferences for several visual features, including motion directions/axes, contrasts, and spatial and temporal frequencies, indicates that the convergence of RGCs onto a single LIN is not random across multiple visual feature dimensions. Consistent with this idea, a previous study revealed that horizontal direction selectivity in LINs was abolished after genetically disrupting horizontal direction selectivity in RGCs^50^.

#### Different Spatial Yet Similar Directional Preferences of Dendrites Indicate Distinct Yet Coordinated Retinal Inputs

Dendrites from the same *DS* LINs prefer distinct spatial locations aligning with retinotopic map of LIN populations (Figure 3D-G) while sharing coarsely similar motion preferences (Figure 5E). This suggests that retinal inputs onto a single LIN follow the retinotopic progression and are coordinated in motion preferences (Figure 7A). Retinotopic progression along dendritic arbors was also reported in tectal neurons of *Xenopus* tadpoles using local stationary stimuli^64^. Our results go beyond that finding by revealing selective input coordination for similar motion direction preferences, even if these inputs are physically distant along the large LIN dendritic arbor.

**Figure 7.**
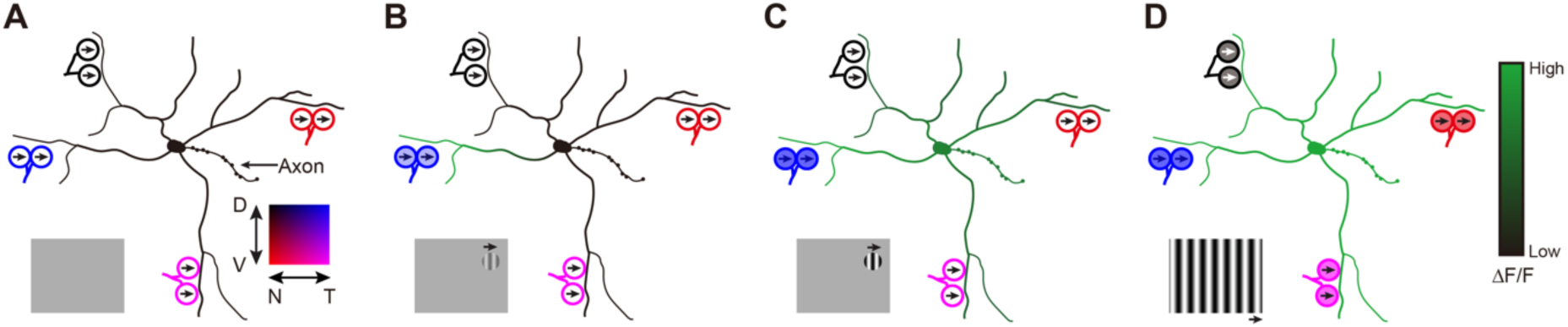
**Local and Global Activation of LINs by Different Visual Stimuli** (A-D) Schematic illustrating distinct activity patterns in one LIN under different visual stimuli. Distinct dendritic compartments of a *DS* LIN are innervated by *DS* RGC axons that are tuned to similar motion directions (indicated by arrows) but different spatial locations (indicated by colors). (A) Under the gray screen condition, no RGC inputs are activated and the LIN remains silent (indicated by black). (B) When local low-contrast drifting gratings are present along the LIN’s preferred direction, a small subset of RGC input is activated, resulting in local activation of the postsynaptic dendrite (indicated by green). The activity propagates along the dendrite and attenuates before reaching the soma. (C) Local high-contrast drifting gratings strongly activate a small subset of RGC input and their postsynaptic dendrite. A large fraction of dendrites of the LIN is activated, with the activity decaying from the dendrite receiving direct RGC input. (D) Strong global activity is evoked in the LIN when many RGC inputs are activated by full-screen high-contrast drifting gratings.

Similar feature preferences among local inputs enable consistent information encoding under global activation. Global activation then results in additional calcium elevation in dendrites on top of their local activation while maintaining or even sharpening the visual feature preference of their local inputs, ensuring that dendritic outputs remain closely matched with the presynaptic retinal inputs. This input coordination distinguishes LINs from parvalbumin (PV)+ interneurons in the visual cortex, which pool inputs with different orientation proferences^65,66^ and provide a general inhibitory tone to their target pyramidal neurons^4^. While random sampling of orientation-selective inputs allows PV+ neurons to perform feature normalization^67^, LINs may convey feature preferences consistent with retinal inputs to TCs via dendrodendritic synapses.

### Size-dependent Direction Selectivity and Response Magnitudes to Motion in *DS* LINs

*DS* LINs showed greater direction selectivity under full-screen stimuli compared to local stimuli (Figures 5 and S4). This size-dependent direction selectivity was only partially inherited from the retina (Figure S5) -- unlike RGC boutons, responses to the preferred direction in LINs increased with larger stimulus sizes. This could be due to nonlinear computation within the LIN: strong local inputs in response to preferred stimuli summing at the soma, triggering action potentials, and evoking global calcium elevation from active dendritic spike propargation^45^. Interaction with other non-retinal inputs could also contribute to this effect. This enhanced inhibition under global stimuli strengthens the inhibition on TCs, potentially suppressing their responses to global motion more effectively and turning the dLGN towards processing local salient stimuli rather than global motion information.

### Possible Roles of Local and Global Responses in LINs

LINs are suggested to control the temporal precision of TC firing by providing short-latency, time-locked feedforward inhibition onto a subset of TCs via local RGCèLINèTC triad synapses, where an RGC terminal synapses onto both a LIN and a TC dendrite and the LIN dendrite also makes a dendrodendritic synapse onto the TC dendrite^16,28^. Our recordings showed that local dendritic activation could happen when low-salience local stimuli are presented at the peripheral receptive fields of the LIN (Figures 4 and S3; see also Figure 7B). Under such circumstances, local dendrodendritic synapses are likely engaged to provide temporally precise local inhibition. Furthermore, the orderly retinotopic organization of retinal inputs along LIN dendrites enables local activation and GABA release from LIN dendrites to be retinotopically matched with their target TCs, thereby providing spatially aligned local inhibition.

When local visual stimuli became more salient, local activation propagated to the soma and evoked global activity that spread to a large part of the dendritic arbor, including dendrites distal to the initiating compartment (Figures 4 and S3; see also Figure 7C). Such global activation likely results in widespread inhibition of TCs across a large region of the dLGN, including TCs responding to visual space far from the stimulus location. This mode of activation may leverage advanced onset inhibition, allowing LINs to exert effective inhibition under salient moving stimuli by providing inhibition before the stimulus even reaches the receptive fields of distal TCs.

A full-screen (global) stimulus could activate all RGC inputs, leading to highly synchronized global responses throughout the dendritic arbors of LINs (Figures 2 and 7D)^45^. In this scenario, LINs could potentially perform spatial normalization^67^ to refine the receptive fields of TCs^68–70^. In the feline dLGN, extraclassical surround suppression is significantly stronger for LGN neurons than for their direct retinal inputs. Moreover, the efficacy of retinal spikes to evoke LGN responses decreases as stimulus size increases, and the onset of suppression is typically fast (∼7 ms delay)^42^. The size-dependent response magnitudes (Figures 5D, 5E and S4E) and temporal characteristics of LINs would make them likely candidates for providing the extraclassical suppression and influencing both size tuning and temporal patterns in TCs.

### Perspectives and Future Directions

To characterize dendritic calcium responses, we focused on direction-selective (*DS*) LINs, as these LINs exhibited significant responses to the visual stimulation set used in this study and had a large portion of their dendrites captured in the same imaging plane, parallel to the surface of the thalamus (we captured 10 such LINs out of 8 mice among 34 total mice we performed *in vivo* imaging). Distinct subtypes of retinal ganglion cells have axon terminals occupying different depths in the dLGN, with *DS* RGCs terminating dorsally in the shell region and alpha RGCs terminating ventrally in the dLGN core^55,71^. Because *DS* LINs sample similar types of inputs across their dendrites, it is logical that they extend their dendrites horizontally in the shell region. Interestingly, a previously reconstructed LIN from electron microscopy (EM) volume extended its dendrites across the entire depth of the dLGN, spanning both the shell and core regions, and was thus speculated to receive mixed visual information across its dendritic arbor^28^. On the other hand, non-*DS* RGC axonal boutons are abundant in the dorsal dLGN region^51^ (see also Figure S1C), suggesting that vertically extended LINs might still receive similar types of non-DS inputs throughout their dendrites. To determine whether the coordination of local and global dendritic activity observed in *DS* LINs also applies to various types of LINs^28,53,72^, new imaging approaches will be needed to record dendrites in the deeper dLGN and rapidly through an extensive range of imaging depth.

In the presence of global signals, dendritic calcium signals could arise from both local synaptic inputs and backpropagating spikes. Quantitatively assessing the contribution of local activation to dendritic calcium signals will provide further insights into signal computation within LINs. For example, it could reveal how distinct dendritic compartments may differentially participate in local dendrodendritic release without triggering global signals and identify the conditions that trigger global responses. Although it is challenging to address this question by direct dendritic calcium imaging, biophysical models of individual LINs could be developed to incorporate their dendritic visual responses measured *in vivo* and dendritic morphology of the same neurons from *post hoc* reconstruction.

Mixed pre- and post-synaptic release sites in dendrites enable neurons to perform input-output computation while minimizing the wiring cost. They are found not only in inhibitory interneurons but also in excitatory neurons, such as mitral cells in the mammalian olfactory system, where backpropagating action potentials synchronize calcium activity across different compartments^73^. This suggests the link between local and global activation, as well as dendrodendritic releases, play important roles in both inhibitory and excitatory computation. Future studies can further elucidate the contributions of this dual-mode computation to information processing across various neuron types and sensory modalities.

## Acknowledgments

We thank members of the Liang and Crair labs for their discussions. Support was provided by the National Institutes of Health (NIH) (R01 EY034697 to L.L.), grants from the Smith Family Foundation, the Whitehall Foundation, the E. Matilda Ziegler Foundation, the Klingenstein-Simons Fellowship Award in Neuroscience, and the Lawrence Young Investigator program (to L.L.), Yale College Dean’s Research Fellowship and Benjamin Franklin College Richter Fellowship (to M.L.).

## Author contributions

LL designed the experiments; LL, YF, Chen L, Chang L, WH, YC, ML, and DZ performed surgeries; LL and KW performed *in vivo* imaging; KW, WH, and LL analyzed data; Chen L, KW, and YC performed histology and neurite tracing; KW and LL wrote the manuscript with inputs from all authors.

---

**Figure S1.**
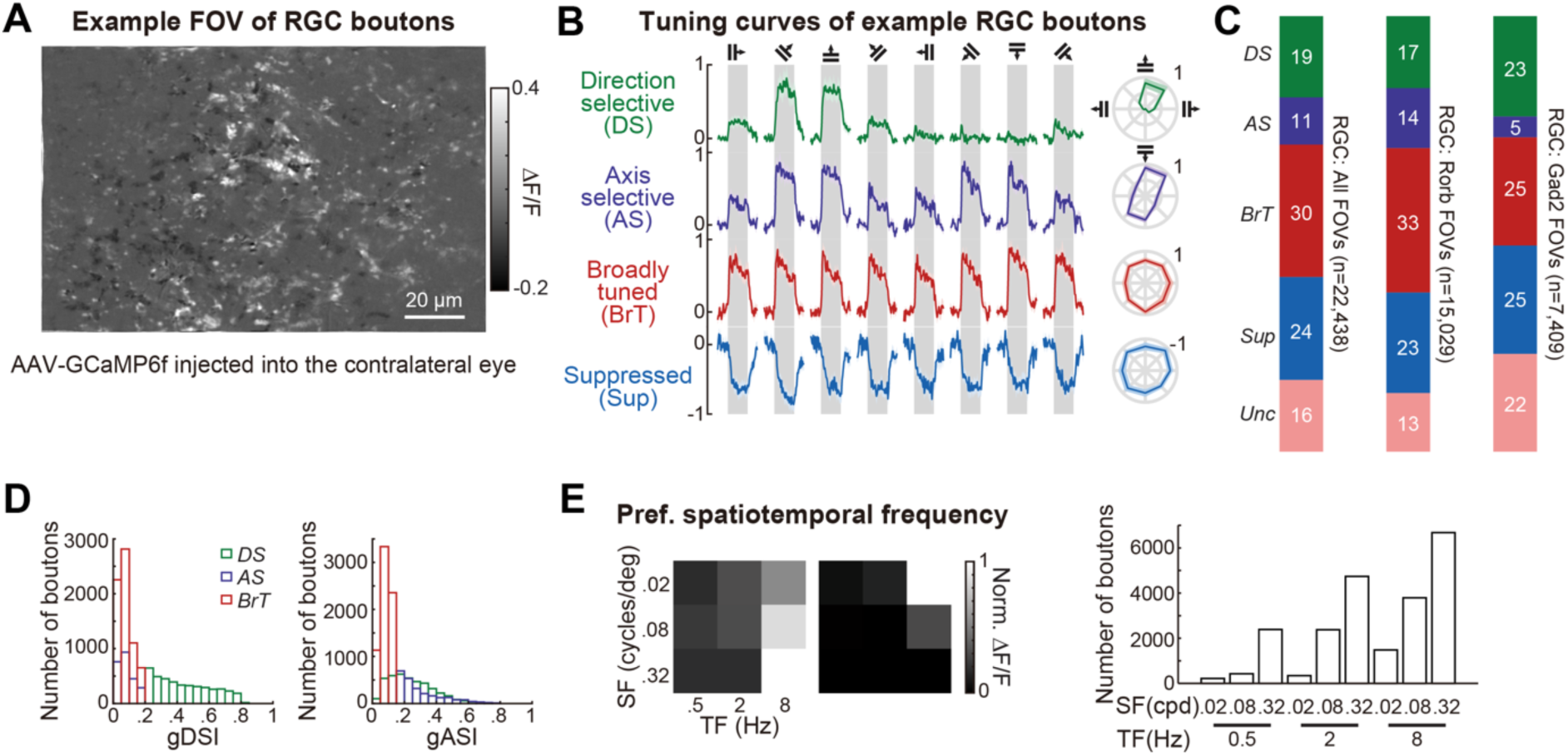
Functional Diversity of RGC Boutons in Response to Motion, Related to. **Figure 1** (A) Visually evoked responses in retinal axonal boutons in an example FOV in the dLGN. To record visual responses of RGC boutons, we injected AAV-CAG-GCaMP6f into the right eye of the mouse and recorded visually evoked calcium signals from their axons in the left dLGN. (B) *Left*, example response timecourses (mean ± SEM) for one example RGC bouton in each functional category (gray bars: 2s presentation of full-screen drifting gratings). *Right*, normalized mean response tuning curves of the example bouton on the left. (C) Proportion of RGC boutons in each category (N = 22,438 responsive boutons from 26 FOV, 11 mice; Rorb-Cre: 18 FOV, 8 mice; Gad2-Cre: 8 FOV, 3 mice). (D) Histogram of gDSI and gASI for *DS*, *AS* and *BrT* boutons. (E) *Left*, response heatmaps of two example RGC boutons preferring different spatial and temporal frequency combinations. *Right*, distribution of the preferred spatial and temporal frequency combinations across RGC boutons (N = 22,438 responsive boutons).

**Figure S2.**
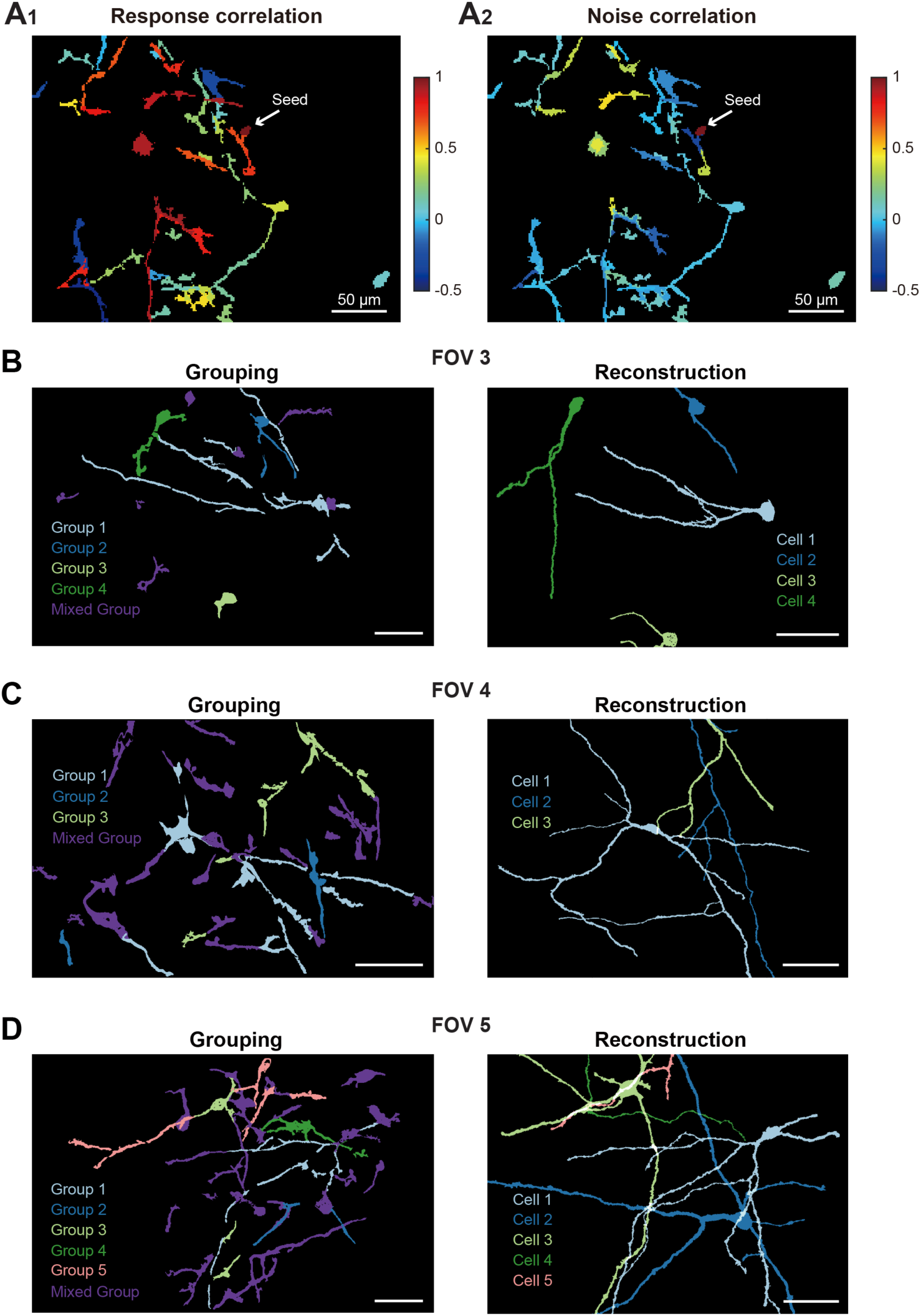
Noise Correlation Is a Better Indicator of Dendrite Identity than Response Correlation, Related to. **Figure 2** (A) Heatmaps of response correlations (A_1_) and noise correlations (A_2_) between a seed dendritic segment and all other segments. Note that the seed segment had high response correlations with segments from different neurons (A_1_) but much lower noise correlations with those segments. (B-D) Grouping results based on noise correlations closely matched the reconstruction of neurons using confocal stacks.

**Figure S3.**
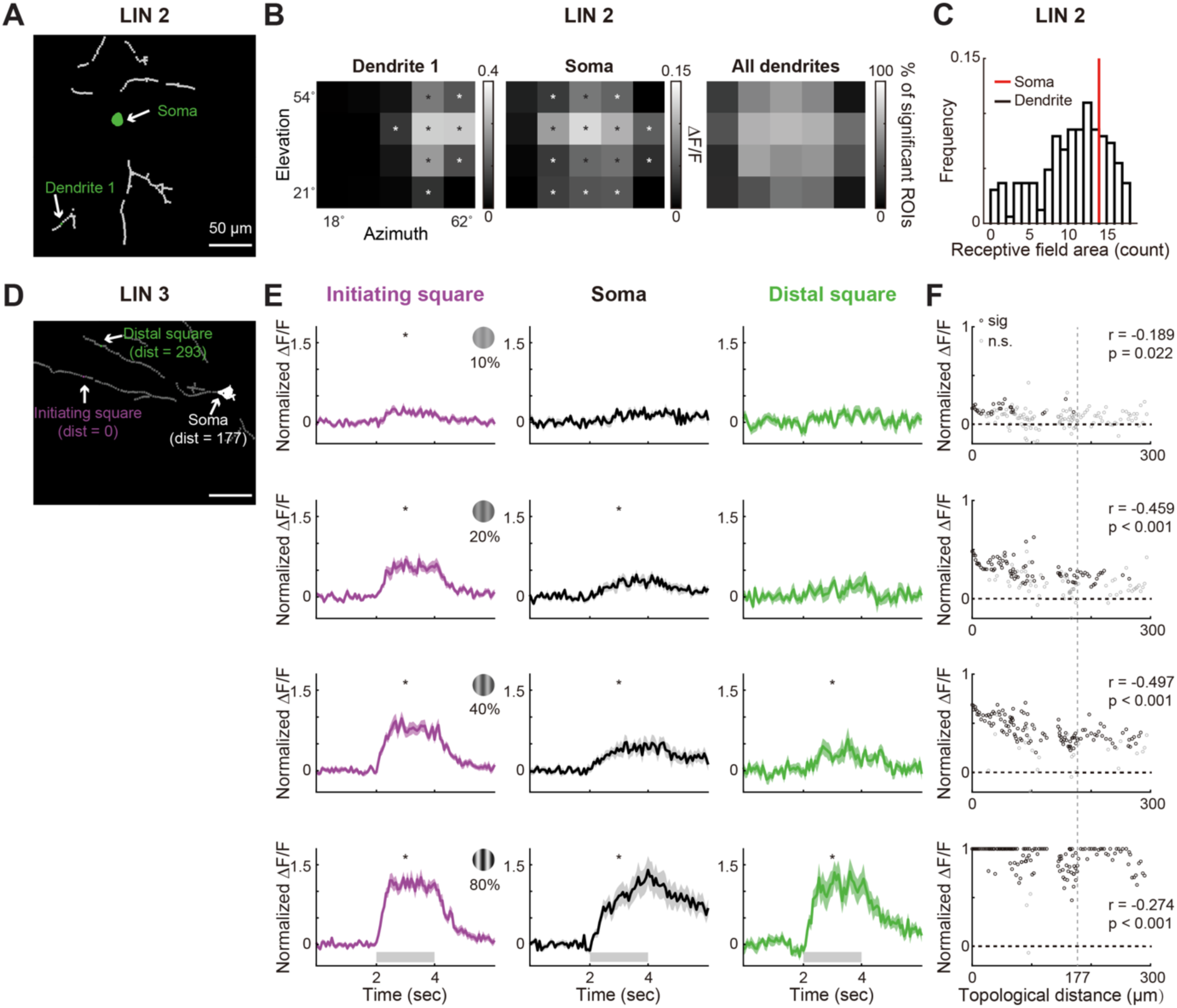
Receptive Fields of the Soma and Dendrites in LIN 2, Related to. **Figure 4** (A) Masks of a second example LIN (LIN 2, the same example neuron in Figure 3) with the soma and an example dendritic ROI highlighted. (B) Visual responses to patch drifting gratings presented at 5 azimuth x 4 elevation locations from the example dendritic square (*left*) and the soma (*middle*). Asterisks (*) indicate the stimulus positions evoking significantly positive and reliable responses from dendrite 1 (*left*) or the soma (*middle*). *Right*, the fraction of dendritic squares from LIN 2 that exhibited significant visual responses to the patch stimuli presented at a given location. (C) Receptive field areas (counts) of the majority of dendritic square ROIs were smaller than that of the soma. Receptive field areas (counts) were calculated as the number of stimulation locations where patch stimuli evoked significant responses from the ROI. (D) Masks of a third LIN (LIN 3) with the initiating dendritic square ROI, a distal dendritic square ROI from another major branch, and the soma highlighted. (E) Average normalized response traces of the ROIs in LIN 3 to patch drifting gratings of 10% (*1st row*), 20% (*2nd row*), 40% (*3rd row*), and 80% (*4th row*) contrasts (mean ± SEM). Asterisks (*) indicate significantly positive and reliable responses. Under low contrast (10%), calcium signals were observed locally but failed to propagate to the soma. With increasing contrasts, calcium signals propagated to the soma and triggered global responses. (F) Scatter plot of normalized responses from ROIs in LIN 3 against their topological distances from the initiating square under patch stimuli with contrasts of 10% (*1st row*), 20% (*2nd row*), 40% (*3rd row*), and 80% (*4th row*). With increasing contrast, response magnitude increased, and significant calcium signals propagated further down the dendrites. Normalized responses and topological distances were negatively correlated under all contrasts (10%: r = −0.189, p = 0.022; 20% contrasts: r = −0.459, p < 10^-8^; 40% contrasts: r = −0.497, p < 10^-9^; 80% contrasts: r = −0.274, p < 10^-3^), suggesting calcium signals propagated with decay.

**Figure S4.**
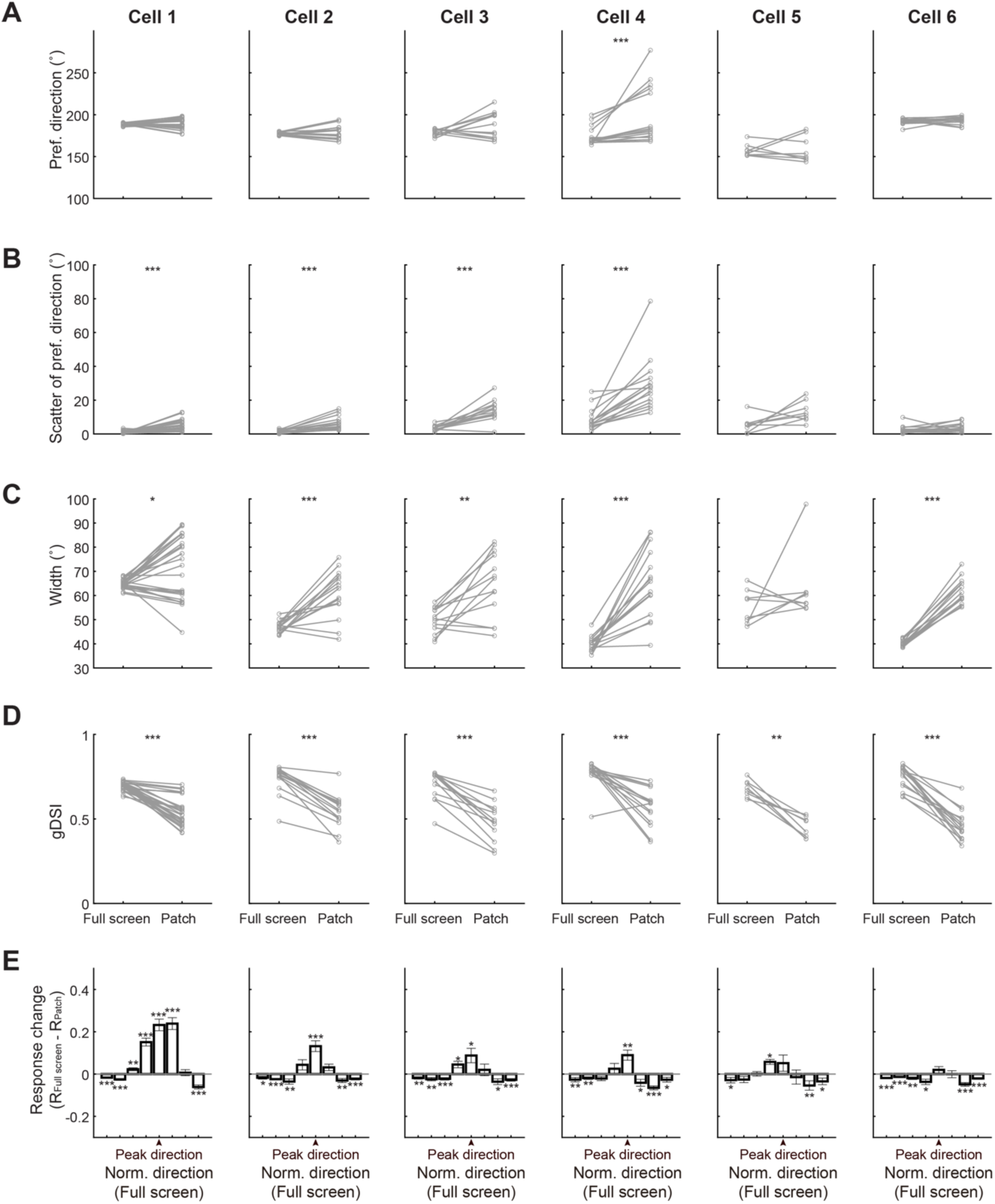
Visual Responses of LIN Dendrites from Individual Neurons to Local and Full-screen Stimuli, Related to. **Figure 5** (A) Preferred directions of dendritic segments under full-screen and patch drifting gratings from individual LINs. No significant changes in preferred directions between full-screen and patch drifting gratings were observed for 5 out of 6 neurons (cell 1, p = 0.219; cell 2, p = 0.421; cell 3, p = 0.110; cell 4: p < 10^-4^; cell 5: 0.844; cell 6, p = 0.277). (B) Scatter of preferred directions (calculated as the absolute differences between the preferred directions of individual dendritic segments and the mean preferred direction across all dendritic segments of the same LIN) under full-screen and patch drifting gratings from individual LINs. 4 out of 6 neurons showed significantly smaller scatter under full-screen than patch drifting gratings (cell 1, p < 10^-4^; cell 2, p < 10^-3^; cell 3, p < 10^-3^; cell 4, p < 10^-4^; cell 5, p = 0.078; cell 6, p = 0.277). (C) Tuning widths of dendritic segments under full-screen and patch drifting gratings from individual LINs. 5 out of 6 neurons showed significantly smaller tuning widths under full-screen than patch drifting gratings (cell 1, p = 0.023; cell 2, p < 10^-3^; cell 3, p = 0.003; cell 4, p < 10^-4^; cell 5, p = 0.388; cell 6, p < 10^-4^). (D) Direction selectivity (gDSI) of dendritic segments under full-screen and patch drifting gratings from individual LINs. All neurons showed significantly higher gDSI under full-screen than patch drifting gratings (cell 1, p < 10^-5^; cell 2, p < 10^-4^; cell 3, p < 10^-3^; cell 4, p < 10^-3^; cell 5, p = 0.008; cell 6, p < 10^-4^). (E) Differences in response magnitudes between full-screen and patch drifting gratings at each of the 8 directions from individual LINs. Responses to null directions were consistently suppressed for all neurons, while responses to preferred directions were enhanced for 4 out of 6 neurons under full-screen drifting gratings. Only *DS* segments under patch and full-screen gratings were included. Wilcoxon signed-rank test was used for individual neuron results in A-E; * p < 0.05, **p < 0.01, ***p < 0.001.

**Figure S5.**
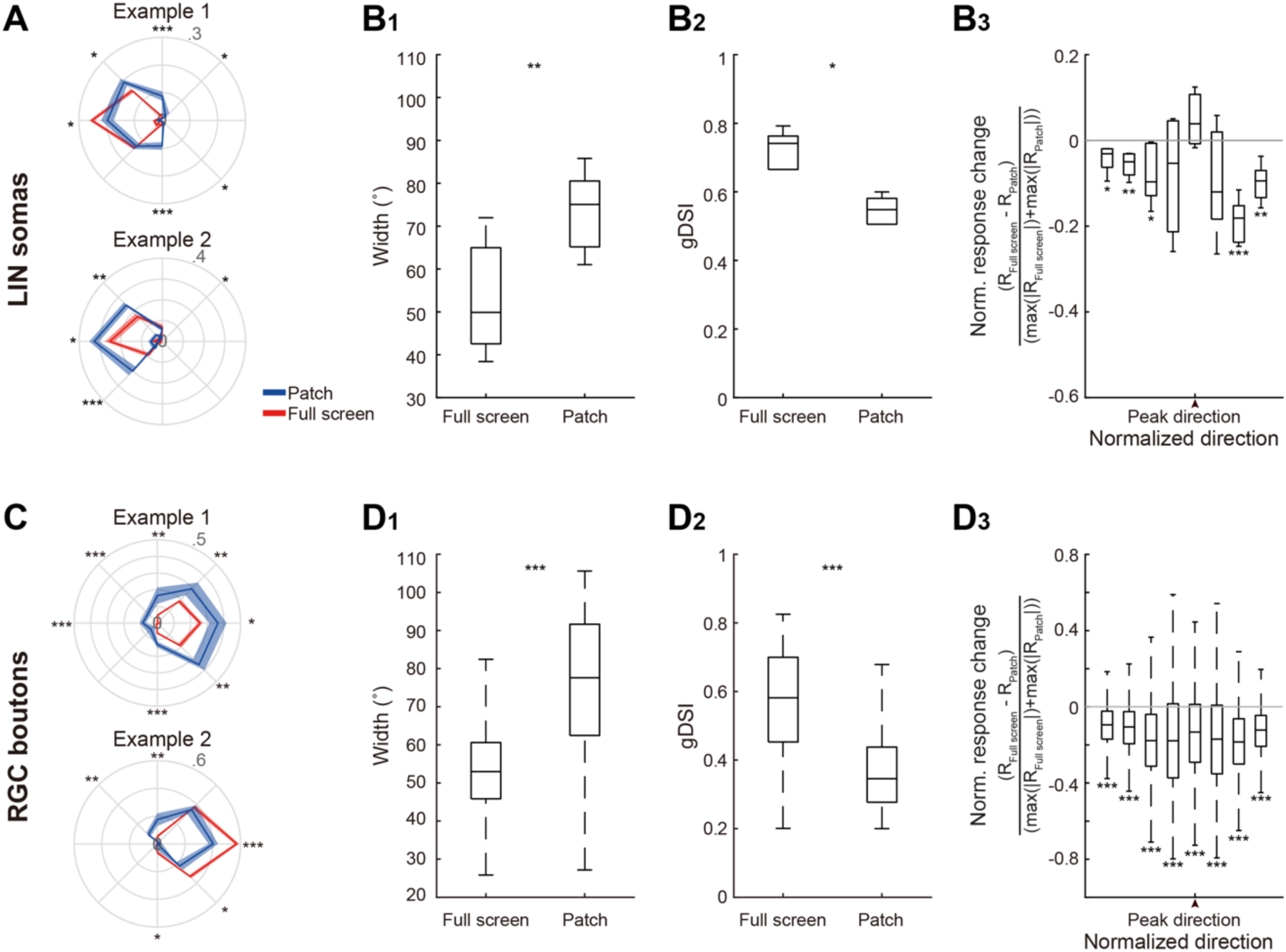
Visual Responses of LIN Somas and RGC Boutons to Local and Full-screen Stimuli, Related to. **Figure 5** (A) Example tuning curves of LIN somas to full-screen and patch drifting gratings. Small negative values are displayed in the radial direction opposite to the measured direction. (B) Box plots of visual response properties of LIN somas to full-screen and patch drifting gratings (B_1_, tuning width; B_2_, gDSI; B_3_, normalized differences in response magnitudes to full-screen and patch drifting gratings). LIN somas showed decreased tuning widths (p = 0.006) and increased gDSI (p = 0.011) under full-screen stimuli. Normalized response changes at null directions were suppressed with increased stimulus sizes (peak direction under full-screen stimuli, p = 0.083; −45°, p = 0.164; −90°, p = 0.020; −135°, p = 0.003; −180°, p = 0.012; +45°, p = 0.078; +90°, p < 10^-3^; +135°, p = 0.002). Linear mixed-effect model was used to control the effects of FOVs. N = 6 somas from 6 FOV, 6 mice. (C) Example tuning curves of RGC boutons to full-screen and patch drifting gratings. (D) Box plots of visual response properties of RGC boutons to full-screen and patch drifting gratings (D_1_, tuning width; D_2_, gDSI; D_3_, normalized differences in response magnitudes between full-screen and patch drifting gratings). RGC boutons showed decreased tuning width (p < 10^-11^) and increased gDSI (p < 10^-84^) with increased stimulus sizes. Normalized response changes at all directions were significantly suppressed with increased stimulus sizes (peak direction under full-screen stimuli, p < 10^-3^; −45°, p < 10^-3^; −90°, p < 10^-4^; −135°, p < 10^-13^; −180°, p < 10^-22^; +45°, p < 10^-7^; +90°, p < 10^-28^; +135°, p < 10^-133^). * p < 0.05, **p < 0.01, ***p < 0.001. Linear mixed-effect model was used to control the effects of FOVs. N = 991 boutons from 10 FOV, 3 mice. Note that somas and boutons were asked to be both significantly direction-selective (*DS*) under patch and full-screen stimuli. All box plots: central mark, median; the bottom and top edges, the 25th and 75th percentiles, respectively; whiskers, lowest and highest data points, excluding outliers.

**Figure S6.**
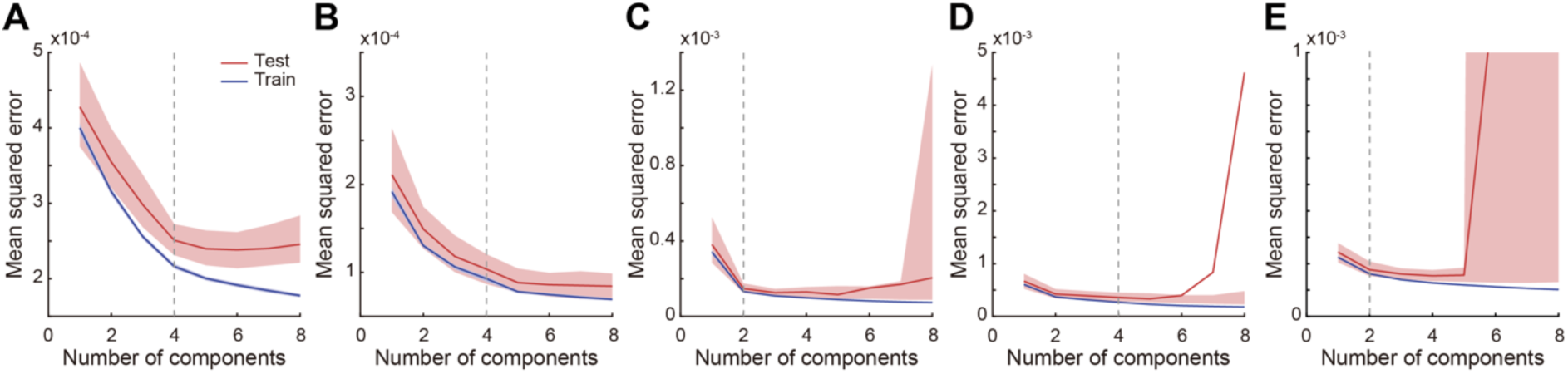
Factorization of Dendritic Responses of LINs, Related to. **Figure 6** (A-E) Mean squared error (MSE) versus the number of model components for training (blue) and test (red) datasets in cross-validation for non-negative matrix factorization of dendritic responses in five additional individual LINs (N_repeats_ = 200; lines, mean; shaded areas, the 95% confidence intervals [CI_95%_]; dashed line, the location of the test MSE closest to 5% above its minimal value). Note that the mean of test data is outside the CI_95%_ in D due to outliers.

## METHODS

### CONTACT FOR REAGENT AND RESOURCE SHARING

Further information and requests for resources and reagents should be directed to, and will be fulfilled by, the Lead Contact, Liang Liang.

### EXPERIMENTAL MODEL AND SUBJECT DETAILS

#### Animals

Gad2-IRES-Cre (JAX: 028867) and Rorb-IRES-Cre (JAX: 023526) were purchased from the Jackson Laboratory. All animal procedures followed the Yale Institutional Animal Care and Use Committee (IACUC) and federal guidelines. Animals were housed on a 12-hour light-dark cycle, with imaging experiments on the same animal conducted across multiple days during the same period each day. Animals were housed with standard mouse chow and water provided ad libitum. Both male and female mice were used in all studies.

### METHOD DETAILS

#### Virus injections

To monitor calcium activity in LINs, a total of 100-200 nL of Cre-dependent AAV encoding the calcium indicator GCaMP6f (AAV2/1.hSyn.Flex.GCaMP6f.WPRE.SV40, Addgene #100837-AAV1; or AAV2/9.CAG.Flex.GCaMP6f.WPRE.WV40, AddGene #100835-AAV9) was stereotaxically injected into the left dLGN in 8∼12-week-old Gad2-IRES-Cre mice. Mice were anesthetized with isoflurane in 100% O_2_ (induction, 3%–5%; maintenance, 1%–2%) and placed on a heating pad (CWE) in a stereotaxic apparatus (KOPF). Ophthalmic ointment (Vetropolycin) was applied to their eyes and extended-release buprenorphine (3.25 mg/kg, s.c.) was administered. Subcutaneous infiltration with lidocaine (less than 7 mg/kg) was applied five minutes before making the skin incision. Two injections were made at 2.25-2.3 mm lateral and 2.3-2.75 mm posterior to Bregma, and 2.6-2.8 mm ventral from the skull, respectively. After surgery, mice were administered meloxicam (0.5 mg/kg i.p.) and monitored during recovery.

To measure visual responses from retinal ganglion cell axons, 1 µl of AAV2/2.CAG.GCaMP6f.WPRE.SV40 (Addgene #100836; packaged by Yale Vision Core) was gently injected intravitreally into the right eye of Gad2-IRES-Cre or Rorb-IRES-Cre mice (both on C57BL/6J background) after the animal was anesthetized by intraperitoneal injection of a mixture of ketamine (100 mg/kg) and xylazine (10 mg/kg). Care was taken to prevent cataract formation during the injection procedure. Mice were administered meloxicam (0.5 mg/kg i.p.) at the end of the procedure and monitored during recovery.

#### Headpost and cranial window implant for dLGN imaging

1-3 weeks after viral injection, a headpost and cranial window were implanted using methods detailed in Liang et al^51^. Mice were administrated dexamethasone sodium phosphate (4.8 mg/kg) approximately 2 hours prior to surgery to reduce brain edema. Mice were anesthetized with isoflurane in 100% O_2_ (induction, 3%–5%; maintenance, 1%–2%) and placed on a heating pad (CWE) in a stereotaxic apparatus (KOPF). Ophthalmic ointment (Vetropolycin) was applied to their eyes, and extended-release buprenorphine (3.25 mg/kg, s.c.) was administered. Subcutaneous infiltration with lidocaine (less than 7 mg/kg) was applied five minutes before making the skin incision. A two-pronged headpost was attached to the skull, centered approximately 2.7 mm lateral and 1.9-2.1 mm posterior to Bregma over the left hemisphere and oriented tangentially to the curved skull surface. The head was then tilted to secure the headpost in clamps that aligned the headpost parallel to the stereotaxic apparatus platform. A 3-mm diameter craniotomy was performed at the center of the headpost. Cortical and hippocampal tissue underneath was gently aspirated to reach the thalamus surface while preserving the thalamic surface and optic tract. A 3 mm x 3.4 mm (diameter x height) stainless steel cylindrical cannula (MicroGroup) with a 3-mm diameter coverslip glued to the bottom by UV-cured Norland Optical Adhesive 71 was stereotaxically inserted into the craniotomy and lowered approximately 2.75 mm below the skull’s surface to slightly press against the thalamus surface. The cannula was affixed to the skull with C&B Metabond (Parkell). To facilitate imaging with a water-immersion objective and light shielding, a low-profile adaptor was created by gluing a neodymium ring magnet (Indigo® Instruments, outer diameter, inner diameter, height: 7.5 mm, 5 mm, 1 mm) to the skull around the cannula. During two-photon imaging sessions, this ring magnet held the light shield in place by contacting an 8 mm x 0.3 mm (diameter x height) spring steel round shim (McMaster) attached to the blackout fabric (Thorlabs). Finally, meloxicam (0.5 mg/kg, i.p.) was administered, and the mouse was monitored during recovery. Mice were given seven days to two weeks to recover before *in vivo* imaging.

#### Epifluorescence and two-photon calcium imaging

To identify the dLGN in the thalamus, we performed epifluorescence imaging (Neurolabware) in head-fixed mice to measure bulk calcium signals in response to visual stimulation displayed at 3 x 3 locations on the monitor. A blue LED light source (470 nm center, 40 nm band, Chroma) was used for excitation, and the green fluorescence was passed through a 500 nm long-pass emission filter and collected using an sCMOS camera (PCO). The light shield around the objective blocked light emitted by the LCD screen. The light was focused on the surface of the thalamus. Epifluorescence images (512 x 384 pixels) were acquired at 15.5 Hz, using a 4x, 0.28 NA objective (Olympus).

Two-photon calcium imaging was performed using a resonant-scanning two-photon microscope (Neurolabware). All images were acquired using a 20x, 1.0 NA, 5.6 mm WD objective (Zeiss) at 2.0x (∼417 x 313 µm^2^), 2.8x (∼298 x 224 µm^2^) or 3.4x (∼253 x 187 µm^2^) digital zoom. The light shield around the objective blocked light emitted from the LCD screen. We focused on imaging fields of view (FOV) at depths of 56-150 µm below the surface of the thalamus (LIN: 56∼108 µm; RGC: 78∼150 µm), using an Insight X3 laser (80 MHz; Spectra-Physics) at 960 nm. Laser power measured at the front aperture of the objective was 30-65 mW – likely a substantial overestimate of the actual power reaching the sample via the cannula. Emitted signals were detected by a GaAsP PMT after a 562 nm longpass dichroic mirror and a 510/84 bandpass filter. Images were collected at 15.6 frames/s, 512 x 796 pixels/frame, using Scanbox (Neurolabware). Each imaging run lasted approximately 30 minutes, and ∼6 runs were typically performed during each imaging session. One FOV was imaged during one imaging session. To measure visual response properties from RGC axons or LIN somas, several different FOVs were typically imaged in each mouse, with each FOV at least 20 µm above or below other FOVs.

The same FOV was imaged in multiple two-photon imaging sessions over approximately two weeks to record responses from dendrites of individual LINs. To facilitate consistent positioning across imaging sessions, we marked the two prongs of the headpost at their contact points with the clamps after the first imaging session and used these marks to guide the positioning of the headpost in subsequent sessions. During two-photon imaging, we first coarsely identified the x and y coordinates of the FOV using a reference image of the superficial thalamus containing second-harmonic generation signals and then the z coordinate using a reference z stack. Fine alignment of the FOV was achieved by carefully comparing it to a reference image from the first imaging session.

Epifluorescence and two-photon imaging experiments were typically performed between one week and one month after headpost and cranial window implantation.

#### Visual stimulation

Visual stimuli were generated using Psychtoolbox^74^, and displayed on a luminance-calibrated LCD monitor (Dell, 17”,1024 x 1280 pixels, 60 Hz refresh rate) placed 22 cm from the mouse’s right eye, spanning approximately 6° to 84° along the azimuth (0° representing the center of the binocular zone) and -7° to 59° along the elevation (0° representing the eye level) of the visual space. The mean luminance of the monitor was 24.5 lux (16.4 lux at the eye).

To measure large-scale retinotopic organization using epifluorescence imaging, local 20° Gabor-like circular patches containing square-wave drifting gratings were presented at 9 retinotopic locations for 2 seconds (drifting along the 180°, temporal-to-nasal direction; 0.08 cycles per degree (cpd); 2 Hz; 80% contrast), followed by 2 seconds of uniform mean luminance, with 20 repeats per stimulus location.

To measure retinotopy progression during two-photon imaging, we used a binarized version of a bandpass-filtered noise stimulus with a spatial frequency corner of 0.05 cpd, a cutoff of 0.32 cpd, and a temporal frequency cutoff of 4 Hz^58^. The noise stimulus was presented within 5° × 40° bars at 80% contrast, displayed vertically at 8 azimuth locations and horizontally at 8 elevation locations, covering an area of 40° × 40°. Stimuli were presented for 2 seconds, followed by a 2-second gray screen at mean luminance during the inter-trial interval. Visual stimulation also included a blank condition (mean luminance). Stimulus order was randomized within a single repeat (consisting of a single presentation of each stimulus condition). The noise bar stimulation was presented for 20 repeats during one run and typically one run per imaging session.

To measure the direction, axis, spatial, and temporal frequency preferences of RGC boutons and LINs, we presented full-screen sine-wave drifting gratings at 80% contrast during two-photon imaging. These gratings were shown in eight different directions (spaced 45° apart) at spatial frequencies of 0.02, 0.08, and 0.32 cpd and temporal frequencies of 0.5, 2, and 8 Hz, respectively. The visual stimulation paradigm also included periods of full-screen mean luminance (gray, blank trials). All stimuli were displayed for 2 seconds with 2-second inter-trial intervals of mean luminance gray. A single repeat consisted of one presentation of each combination of the direction, spatial frequency, and temporal frequency (72 combinations), interleaved with 9 presentations of ‘blank’ stimuli, all in random order. Typically, 12∼30 repeats were recorded per imaging session, each with a different randomization of the condition order. To measure the visual response properties of individual LINs or RGC boutons to local stimuli, we presented 10° or 20° (10° only for RGC boutons) Gabor-like circular patches containing sine-wave drifting gratings at 9∼25 retinotopic locations (12 for RGC boutons). These gratings were delivered under the preferred spatial and temporal frequency combination for individual LINs or 0.08 cpd, 2 Hz for RGC boutons. The gratings were presented in the preferred and null directions for one LIN, and in all eight directions (spaced 45° apart) for all other LINs and RGC boutons, for 2 seconds. The visual stimulation paradigm also included periods of mean-luminance gray (gray, blank trials). The inter-trial interval (mean-luminance gray) lasted 2 seconds for 10° patches and 3 seconds for 20° patches. A single repeat of visual stimuli involved one presentation of each direction at each location, and 9∼25 presentations of ‘blank’ stimuli, in random order. Typically, 4-6 repeats were presented during one run, and 3∼4 runs of local stimuli per imaging session, each with a different randomization of trial orders.

To compare visual responses of the same LINs or RGC boutons under local vs. full-screen stimuli, we also presented full-screen sine-wave drifting gratings in 8 directions under the same spatial and temporal frequency used for local stimuli. Stimuli were displayed for 2 seconds, with 4-second mean-luminance gray during inter-trial intervals. A single repeat involved one presentation of each direction and one presentation of ‘blank’ stimuli, interleaved in random order. A single run of full-screen stimuli consisted of 30-50 repeats. One full-screen run was usually presented in the middle of the imaging session, with runs of local stimulation delivered both before and after it. We also examined visual responses to patch and full-screen stimuli under additional contrasts (10%, 20%, and 40%) in three LINs at their preferred directions.

For imaging LINs in each mouse, we first mapped the retinotopy under noise bar stimulation and full-screen motion preferences during early imaging sessions. Visual response properties under local stimuli and contrast sensitivity were mapped during subsequent imaging sessions.

#### Immunohistochemistry, confocal imaging, and *post hoc* reconstruction

We performed immunohistochemistry, *post hoc* neurite tracing, and reconstruction of individual neurons after *in vivo* two-photon imaging sessions. Mice were terminally anesthetized with Euthanasia III solution (>150 mg/kg) via intraperitoneal injection and transcardially perfused with 1x phosphate-buffered saline (PBS) immediately followed by 10% formalin. Brains were post-fixed overnight at 4°C and subsequently transferred into 20% sucrose solution. Cryosections of 150–200 μm thickness were made using a microtome (Thermo Scientific HM430) tangential to the dLGN. Brain sections were incubated in a primary antibody solution containing 3% donkey serum, 0.4% Triton X, and 0.02% sodium azide in PBS for 3-7 days at 4°C, followed by three washes in 0.4% PBST buffer and incubation with secondary antibodies in 0.4% PBST buffer containing 0.02% sodium azide for 2–5 days at 4°C. After three washes in PBS and one hour clearing in 67% thiodiethanol (TDE, Sigma-Aldrich)^75^, brain sections were mounted in VECTASHIELD Antifade Mounting Medium with DAPI. The primary antibodies used were chicken anti-GFP (1:1000, Invitrogen, A10262; for staining of GCaMP6f). The secondary antibodies used were donkey anti-chicken Alexa Fluor 488 (1:800, Life Technologies, 703-545-155). Tiled z-stack images were collected using an LSM 800 confocal microscope (Zeiss) with a 63x, 1.4 NA objective and stitched using ZEN 3.5 (blue edition). Individual neurons were traced double-blindly using the Simple Neurite Tracer plugin in Fiji^76^.

#### Image preprocessing

To correct for x-y motion in images acquired from two-photon imaging, a series of image registration and denoising steps were applied^51^. For each imaging session, a reference template was generated by averaging the first 500 frames of the first run after one round of translational registration of these 500 frames using efficient subpixel registration methods^77^. The movies taken within the same imaging session were first aligned to the reference template using translational registration. Next, the registered movies were denoised to aid further image registration. Specifically, principal component analysis (PCA) was performed to identify spatial principal components of the field of view, and each image of the movie was reconstructed using only the first 400 (with the highest eigenvalues) out of approximately 30,000 principal components. The denoised images were then subjected to local normalization and image warping to derive the warping parameters, which were subsequently applied to the original raw movie. As a final step, PCA de-noising was performed again to increase the signal-to-noise ratio and facilitate mask extraction of neuronal structures. The observed results were not dependent on this last step. For additional details, please see Liang et al^51^.

#### Bouton mask extraction

Non-overlapping retinal bouton masks were automatically extracted based on average images of absolute change in fluorescence (ΔF/F) using our previously developed algorithm^51^. To remove residual calcium signals not originating from boutons themselves, we estimated neuropil masks as circular annuli of 3 µm width, with the inner edge located 2 µm beyond the edge of a corresponding bouton mask. Pixels from adjacent bouton masks were excluded from these neuropil masks.

#### Soma mask extraction

We used suite2p to extract non-overlapping soma masks for local interneurons^78^. The timescale (in second) used for the deconvolution kernel is 0.7. The sampling rate is 15.6 Hz. Thresholding scaling was set at 0.5 to extract neuron masks. Preprocessed movies from 2 to 4 runs of the same imaging session, but under different visual stimuli, were provided to the algorithm. The resulting spatial footprints were used as soma masks.

#### Dendrite mask extraction

Dendritic regions of interest (ROIs) were detected using a constrained non-negative matrix factorization (CNMF) approach implemented in Python as part of the CaImAn software package^61,62^. We used the ‘sparse_nmf’ initialization method. For each imaging session of each mouse, 2 to 4 preprocessed movies from runs with different visual stimuli were fed to the algorithm. The algorithm returned two matrices of spatial and temporal components, respectively. Each spatial component was reshaped into a (height x width) matrix, with pixel values representing the weight of this pixel to the corresponding temporal component (a 1 x timepoints vector, representing the time course of the spatial component). The first 60 spatial components were kept and further segmented according to their spatial connectivity with the following procedures.

We first cleaned CNMF spatial components with Gaussian blurring and local normalization. Initially, a Gaussian filter (sigma = 2.62 µm) was applied to all spatial components, and pixels were thresholded by a z-score of 1.5. The thresholded components containing neuronal structures were then subjected to a second round of Gaussian filtering (sigma = 2.62 µm) and thresholding with a z-score of 2.5. For components that lost meaningful neuronal structures after the first round of Gaussian blurring and thresholding due to their low weights, we returned to their original CNMF components, applied local normalization (the standard deviations of the 2-D Gaussian smoothing kernel for computing the local mean and local variance were 4.19 and 20.43 µm, respectively), and thresholded the image with a z-score of 0.5.

Next, we binarized each cleaned component and performed the first round of image segmentation within each CNMF component by extracting spatially connected pixels into non-overlapping individual segments using the ‘CellsortSegmentation’ function from CellSort 1.0^60^. To avoid overlaps between segments from different components, we summed all binary segment masks into an overlap matrix, where the values of each pixel indicated the number of segments containing that specific pixel. Occasionally, patchy and smeared neuropil areas survived up to this step, with their pixel values typically being 1. We manually identified and excluded these neuropil pixels by setting their values to 0 using the MATLAB built-in function ‘roipoly’. After this, we isolated each overlap layer (i.e. we binarized the overlap matrix by only keeping pixels with values equal to i, i = 1…max(overlap_matrix)) and further segmented them by pixel spatial connectivity.

To remove background calcium signals not originating from the neuronal structure in LIN FOVs, we estimated neuropil masks as circular annuli of 4 µm width, with the inner edge located 4 µm beyond the outermost edge of a corresponding soma or dendrite mask. Pixels from adjacent soma and dendritic masks were excluded from these neuropil masks.

#### Cross-day alignment of ROIs from the same FOV

To thoroughly explore the response properties of individual LINs, we returned to the same FOV across multiple imaging sessions. We aligned the ROI masks across imaging sessions after preprocessing the movies from each session respectively. For each imaging session, we generated its reference template by averaging the first 500 frames of its first run. We designated one of the image sessions as the reference session and aligned its template to each of the other templates by performing translational registration and warping (using the MATLAB built-in function ‘imregdemons’). Subsequently, we applied the same translational shift and displacement field from the warping to the ROI masks extracted from the reference session to obtain the aligned ROI masks for the other imaging sessions.

#### Time course extraction and correction

To obtain raw fluorescence traces for segment masks and neuropil masks, the fluorescence intensity value of a segment at each time point was defined as the average fluorescence across the pixels belonging to the mask.

To account for neuropil signals that may contaminate signals in the segment trace, we applied neuropil correction by subtracting a scaled version of the corresponding baseline-removed neuropil trace 𝑁(𝑡) from each segment trace 𝐹(𝑡)^79^:

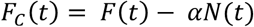

To determine the scale factor α, we used the following procedures first to remove the baseline for segment and neuropil traces and then fit the lowest envelope of the scatterplot for baseline-removed traces of segment versus neuropil. Specifically, we divided the raw trace into equal-sized bins (bin size = 50 frames or 3 s) and populated each bin with the lowest value from that bin. Next, we median-filtered the baseline trace (window = 3000 frames or 192 s) and subtracted that from the raw trace. Subsequently, we binned the baseline-removed neuropil traces (bin size = 800 frames or 51 s), and we identified the frame with the lowest value of the segment trace within each bin. We performed linear regression using these lowest values and determined α as the slope of the regression line. The average α was 0.43; we set the upper and lower bound of α to 1 and 0. A similar procedure was subsequently performed for each of the other segments that had an overlap of at least 1.6 µm^2^ with the neuropil mask for the segment of interest to correct influences from other neuronal structures.

We also corrected baseline fluorescence F_0_ to remove the decay in fluorescence from the activity evoked during the previous visual stimulus presentation using our previously developed algorithm^39,51^. Due to calcium buffering and GCaMP6f buffering, the segment fluorescence sometimes did not fully return to the baseline during the 1 second after the offset of a previous stimulus presentation and persisted in the next second used to calculate F_0_ for the following stimulus period. Therefore, a baseline correction was introduced that modeled this exponential decay during the period of the inter-trial interval from previously evoked GCaMP6f activity. For each segment and each single trial, a fitted, exponentially decaying contribution from the previous trial was subtracted from F(t) during the 1 second before and the 2 seconds during the visual stimulus presentation. The corrected baseline was used as the new baseline F_0_ to compute ΔF/F_0_(t) during the visual stimulus presentation. A single response value during a given trial was obtained by averaging the ΔF/F_0_(t) response during the 2-second stimulus window.

#### Grouping dendritic segments into individual neurons

Neuron identification – the process of assigning dendritic segments imaged in a field of view to the same putative neuron – was carried out on data from full-screen drifting grating runs. Segments with a maximal quality index (QI, the standard deviation of the mean timecourse divided by the mean standard deviation of timecourses from all repeats)^80^ greater than 0.4 across all visual stimulation conditions were considered ‘good’ and retained for further grouping analyses.

Hierarchical clustering based on noise correlation was performed on the retained segments. First, we subtracted the mean ΔF/F traces during the 2-second visual stimulation for a given stimulation condition from single-trial ΔF/F traces of the same condition. These residual single-trial ‘noise’ traces were then concatenated across all conditions into one noise trace for each segment. We focused on periods containing significant fluctuations, as these periods reflected robust noise. To identify such periods, we thresholded each noise trace by 2 standard deviations above or below zero. Additionally, time points within a 700 ms interval before and after each thresholded event were included to capture more complete fluctuation events.

For each pair of dendritic segments, we concatenated the traces for each segment respectively based on the union of the time windows containing significant fluctuation events in either segment. We then computed Pearson correlation coefficients for these two traces. This process was repeated for all pairs of ‘good’ segments in the field of view, resulting in a matrix of pair-wise noise correlation coefficients.

To obtain a sparse matrix populated only by large values, the original matrix was further thresholded: The correlation coefficients for a given segment were set to 0 unless they exceeded 0.5 (coefficient threshold) or 2 standard deviations above the mean values of all the coefficients between that segment and others (standard deviation threshold). The cosine similarity between every pair of segments was then computed from this thresholded matrix. The cosine similarity reflected the cosine of the angle between the vectors of pairwise noise correlation coefficients for each segment.

Next, we grouped segments into clusters using agglomerative hierarchical clustering based on the pairwise distance, computed as ‘1 – cosine similarity’. The cluster distance, representing the distance between two groups of segments, was calculated using the weighted-pair group method with arithmetic means (WPGMA) algorithm. We defined ‘correlation similarity’ as ‘1 – cluster distance’. Segments were assigned to the same neuron if their correlation similarity exceeded a threshold of 0.15. For 3 out of 9 FOVs, a different set of parameters was used due to a different signal-to-noise ratio resulting from increased GCaMP6f expression under the CAG promoter (coefficient threshold: 0.6; standard deviation threshold: 2; correlation similarity threshold: 0.35).

To validate our grouping method, neurites in 5 out of 8 FOVs were traced and reconstructed using confocal image stacks obtained after *in vivo* two-photon imaging. The grouping and reconstruction results showed a high degree of match (Figure 2, F and G; Figure S2, B to D).

#### Significant and reliable responses

An ROI exhibited *significant* responses under a given visual stimulation condition if at least 10 out of 31 frames (15.6 Hz frame rate x 2 second stimulus) from the average trace exceeded 2.5 standard deviations above or below the mean baseline activity (computed using the 1-second window prior to stimulus onset).

An ROI was considered *reliable* under a given visual stimulation condition if its quality index (QI) exceeded 0.3.

### Comparison of the retinotopic progression and receptive fields of dendrites

#### Retinotopic progression under noise bar stimuli

To map the population retinotopy progression of a field of view (FOV), we padded extra pixels on the image edges and divided the FOV into equal-pixel-sized ROIs (5.24 µm x 6.11 µm each). Next, ROIs on the edges of the FOV and pixels from the neuron of interest were removed. For each remaining ROI, we extracted its response time courses to noise bar stimuli. To identify the receptive field center of each ROI, we independently fit its retinotopic tuning curves along the azimuth and elevation axes with Gaussian functions 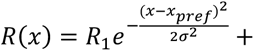 𝑅_offset_. Here, the ΔF/F response 𝑅(𝑥) varies as a function of the noise bar location 𝑥, with the peak response evoked at 𝑥*_pref_*, the preferred retinotopic location. We increased the number of fitting points from 8 to 15, using a previously developed interpolation method^51^.

To determine whether to include an ROI in the retinotopic map calculation, we required the ROI to have significant responses to noise bars and well-fitted retinotopic tuning curves. An ROI was considered to have significant responses if at least 1 out of the 2 bar orientations (azimuth and elevation) had significant responses under at least 2 out of 8 bar locations. The retinotopic tuning curves for an ROI were considered well-fitted if the 𝑟^7^ of fitting (the coefficient of determination) exceeded 0.8, and if the fitted peak amplitude was positive for non-suppressed ROIs or negative for suppressed ROIs (i.e., at least 1 out of 2 bar orientations had significantly negative responses under at least 2 bar locations). ROIs that did not meet these criteria were set to NaN. All the pixels within an ROI were assigned their respective preferred retinotopic locations along the azimuth and elevation axes according to the center location of the Gaussian fit for the ROI. We then convolved the pixel matrix using a Gaussian filter (kernel size = 47 µm, sigma = 10 µm) to generate a smoothed retinotopic map of the entire FOV. To predict the receptive field center for a given dendritic segment according to the population retinotopic progression, we used the pixel value from the retinotopic map at the center of the segment.

#### Receptive field fitting of LIN segments under patch stimuli

To obtain a finer resolution of the receptive field change along dendrites, segments within the same LIN were manually divided into equal-sized dendritic squares (2.62 µm x 3.05 µm) using the ROI Manager in ImageJ. To estimate the receptive field center of dendritic squares under Gabor-like patch drifting gratings, we used conditions under the preferred direction (measured by full-screen gratings) and performed 2D Gaussian fitting of the response magnitudes at different patch locations. The data matrix was linearly interpolated for better estimates. The Gaussian function used was:

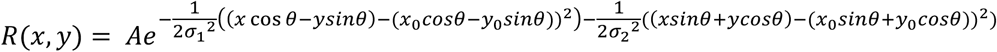

Here, R(x,y) denotes the ΔF/F signal at retinotopic stimulus location x, y; A represents the maximal response evoked at the preferred retinotopic location (x_0_, y_0_); the standard deviation σ_1_ and σ_2_ of the Gaussian are proportional to the receptive field size along the two axes; and θ represents the rotation angle of the 2D Gaussian. We performed positive and negative fitting (restricting A to be non-negative or non-positive) separately and used the fitted result with the larger coefficient of determination (𝑟^7^).

A bootstrapping method involving random sampling of trials from each condition was implemented to fit the retinotopic curves. Specifically, for each of the 100 iterations, we randomly sampled (with replacement) responses from the 6∼18 trials of each condition and fitted the retinotopic curve using the mean responses. The final parameters of the 2D Gaussian were the mean of the fitted parameters across the 100 sampling iterations.

We used a combination of criteria to determine if the procedure yielded a high-quality fit. Each iteration of the fitting procedure yielded a coefficient of determination, 𝑟^7^, defined as the explained variance under least-squares regression. As a second control step, a combined coefficient of determination, 𝑟^2^_*comb*_, was computed by comparing the original retinotopic curve with the fitted curve derived by using the mean of each fitting parameter across the 100 iterations. To assess both the quality and the reliability of the fitting, we used our previously established goodness of fit^51^, 𝐺*_fit_*:

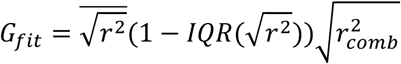

where IQR was defined as the interquartile range – the difference between the 75^th^ percentile and the 25^th^-percentile of the √𝑟^7^ values across iterations. A segment was considered to have a well-fit retinotopic curve in a given bar orientation if 𝐺_6?,_ was greater than 0.7.

### Estimating visual tuning preferences in LINs and RGC boutons

#### Direction tuning curve fitting

A direction tuning curve was computed for each region of interest (including LIN dendrite, soma, or RGC bouton) that showed a significant positive response under at least one direction at a given spatial and temporal frequency combination. The direction tuning curves were initially sampled in steps of 45°. To obtain a more precise estimate of the preferred motion axis and direction, we fitted the direction tuning curves as the sum of two Gaussians^81^:

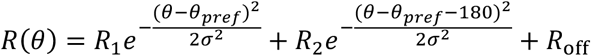

Here, 𝑅(𝜃) is the ΔF/F response for the stimulus direction 𝜃; 𝜃*_pref_* is the preferred direction evoking the peak ΔF/F response 𝑅_1_ ; 𝑅_2_ is the amplitude of the second peak located at 𝜃*_pref_* + 180° ; 𝑅_off_ is a constant amplitude offset; and 𝜎 is the standard deviation. This model assumes that the two Gaussians peak 180° apart and share a common standard deviation.

We implemented several previously reported steps to improve the reliability of the fitting of direction tuning curves and to optimize the accuracy of estimation of the preferred direction of motion^51^, including symmetrically padding the tuning curves to the range of [-90°, 450°] and increasing the number of input points for the fitting procedure from 8 to 25 using a heuristic method of interpolation and extrapolation. We added two shadow copies of the Gaussian function, circularly shifted by +360° and −360°, for the final fitting: 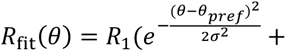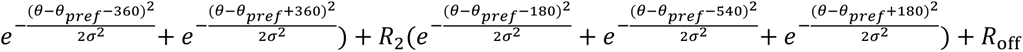

A bootstrapping method was implemented to obtain the fitting parameters and evaluate the reliability of the direction tuning curve fitting. Specifically, we performed 100 iterations of random sampling with replacement from all the trials under each of the 8 directions. For each iteration, we calculated the mean response for each of the 8 directions. These 8-point tuning curves were then interpolated, extended, and finally fitted using the method described above. The final parameters used were the mean of the fitted parameters across the 100 sampling iterations. We required 𝐺_6?,_ to exceed 0.67 to include only ROIs with a high-quality fit (see ‘Receptive field fitting under patch stimuli‘).

#### Axis and direction selectivity

For each region of interest, we calculated a global axis selectivity index (ASI; i.e., selectivity for motion along a given axis) and a global direction selectivity index (DSI) under the spatial and temporal frequency of the peak condition^58,82^, as follows:

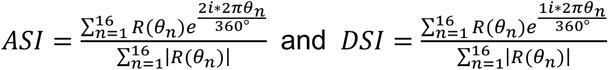

These indices were calculated using the interpolated direction tuning curves, with the ΔF/F response for each of the 16 directions in the range between 0° and 360°. ASI and DSI ranged from 0 (minimum selectivity) to 1 (maximum selectivity). The ASI and DSI computations were iterated 100 times by bootstrapping with replacement from all trials under each of the 8 directions. The final indices were the mean values across the 100 iterations.

### Functional classification of ROIs based on their responses to motion

An ROI was classified as ‘direction-selective’ if for the spatial-temporal frequency combination of the peak condition (by the absolute value of ΔF/F mean) (i) the peak condition was positive, (ii) the positive peak was at least 3 times greater than the negative peak across the 8 directions of stimuli, (iii) significantly positive and reliable responses were observed under at least 1 direction, (iv) the tuning curve was successfully fit (goodness of fit greater than 0.67), (v) the direction selectivity index (DSI) exceeded 0.2 (a value equivalent to 0.33 if direction selectivity was calculated as 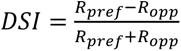, where 𝑅*_pref_* was the response at the preferred direction and 𝑅*_opp_* was the response at the opposite direction).

An ROI was defined as ‘axis-selective’ (i.e., most strongly responsive to motion along two opposite directions constituting a single axis of motion) if for the spatial-temporal frequency combination of the peak condition (by the absolute value of ΔF/F mean) (i) the peak condition was positive, (ii) the positive peak was 3 times greater than the negative peak, (iii) significantly positive and reliable responses were observed under at least 2 directions, (iv) the tuning curve was successfully fit (goodness of fit greater than 0.67), (v) the axis selectivity index (ASI) exceeded 0.15 (a value equivalent to 0.33 if axis selectivity was calculated as 𝐴𝑆𝐼 = 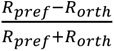, where 𝑅*_ref_* was the response at the preferred direction and 𝑅*_orth_* was the mean response at the two directions orthogonal to the preferred one), and (vi) the DSI was less than 0.2. We distinguished the term axis selective from orientation selective, as the latter is often used even for responses to stationary (i.e., non-drifting) oriented gratings – a stimulus not examined in this study.

An ROI was defined as ‘broadly-tuned’ if for the spatial-temporal frequency combination of the peak condition (by the absolute value of ΔF/F mean) (i) the peak condition was positive, (ii) the positive peak was 3 times greater than the negative peak, (iii) it had significantly positive and reliable responses under at least 3 directions, and (iv) the ASI and DSI were below 0.15 and 0.2, respectively.

An ROI was defined as ‘suppressed’ if (i) the peak condition was negative, (ii) it had significantly negative and reliable responses under at least 3 directions for at least 2 spatial-temporal frequency combinations, (iii) the negative peak was 3 times greater than positive peak under the spatial-temporal frequency combination of the peak condition.

An ROI was defined as ‘unclassified’ if (i) the ROI had significantly positive or negative and reliable responses under at least 3 directions for at least 1 spatial-temporal frequency combination, and (ii) it was not classified as direction-selective, axis-selective, broadly-tuned, or suppressed.

Finally, a small proportion of candidate ROI masks were not significantly driven by any of the presented visual stimuli. These were classified as ‘unresponsive’ and not considered further.

#### Preferred direction or axis of motion

The preferred direction was defined as the fitted 𝜃_5’(6_ values under the spatial and temporal frequency of the peak condition for direction-selective regions of interest. Estimates of the preferred direction ranged from 0° and 360°.

The preferred axis of motion was defined as the fitted estimate of the preferred axis under the spatial and temporal frequency of the peak condition for axis-selective regions of interest. Estimates of the preferred axis of motion ranged from 0° and 180°.

#### Comparison of visual tuning properties between patch and full-screen drifting gratings

To compare the tuning properties of LIN dendrites from the same neuron under patch versus full-screen gratings, we first selected the optimal patch stimulation location for each dendritic segment. For each dendritic segment, we normalized its mean ΔF/F during each 2-sec visual stimulus by its peak mean ΔF/F across all stimulation conditions. We then averaged the normalized mean ΔF/F across segments from the same neuron for each stimulation condition.

A normalized residual for each segment under each stimulation condition was generated by subtracting this average from its normalized response. We then averaged the residuals across directions within each location (‘average residual’) by segment. The location with the largest average residual was identified as the optimal patch location for a given dendritic segment. This method was sensitive to the real preferred location of the segment.

For LIN somas and RGC boutons, the location of the peak condition was considered the optimal patch location. Note that for each LIN soma or RGC bouton, responses to the optimal patch grating were asked to be *DS* for this analysis. Additionally, the optimal patch grating was required to be within 10° of the fitted noise bar receptive field center of that soma or bouton if the responses of the soma or bouton to noise bar stimuli were well-fitted along both the azimuth and elevation axes (see the criteria for a well fit in ‘Retinotopic progression of the FOV under noise bar stimuli’). In cases where noise bar fitting did not meet the criteria, the optimal location was asked to not be on the edge of the stimulation grid (neither the first nor last row or column of all the stimulation locations). Otherwise, the soma or bouton would be excluded.

### Non-Negative Matrix Factorization (NNMF) Model

To extract common features from dendritic responses to patch drifting gratings, we performed non-negative matrix factorization (NNMF) of the dendritic response matrix *A* (see Figure. 6). Each data entry of *A* represented the average ΔF/F response across trials for a dendritic square ROI under a visual stimulation condition (a direction and a location of the patch drifting grating stimuli). Each row of A represented visual responses from one dendritic square ROI across different stimulation conditions, while each column represented responses across different dendritic square ROI under a given visual stimulation condition.

The NNMF decomposed A (dendritic square ROI x condition) into low-rank non-negative matrices 𝑈 and 𝑉^L^, with 𝑉^L^ (component x condition) representing a set of response components that captured the elementary response patterns across visual stimulation conditions and 𝑈 (dendritic square ROI x weight) representing the weights of different response components for dendritic squares. The optimal number of response components was determined using a cross-validation procedure for matrix decomposition, described below^63,83^. In each iteration, we randomly assigned 90% of data entries in A as the training dataset and the remaining 10% as the test dataset, as indicated by a binary matrix M (1 for data used for training and 0 for data used for testing). This resulted in the following optimization problem:

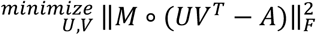

Here, 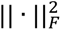 denotes the squared *Frobenius norm* of a matrix, which is the sum of the squares of all its elements.

The optimization problem was resolved by alternately minimizing the above equation for 𝑈 and 𝑉*^T^*. The mean squared error (MSE) of the model was evaluated on the held-out test data points as follows.

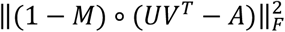

We iterated this procedure 100 times for each given number of response components and calculated the mean and confidence interval of the MSE. We chose the model with 5% more MSE than the minimal MSE to determine the optimal number of components. In this way, we included the components that captured major common response features without losing much information or overfitting. To obtain the final weight and component matrices, NNMF was performed again on the whole data set A with the optimal number of components.

### Detection of the initiation dendritic square

The dendritic square ROI that ‘initiated’ the calcium responses along the dendrites of individual LINs was detected based on dendritic responses to patch stimuli. These stimuli were displayed at the neuron’s preferred direction, spatial and temporal frequency (measured under full-screen stimuli), but at various contrasts (blank, 0.1, 0.2, and 0.4) and locations. For each of the three contrasts (0.1, 0.2, and 0.4) at a given stimulation location, we identified the peak dendritic square (the dendritic square that showed the peak ΔF/F mean) and required it to have significant responses. If at least 2 out of 3 contrast levels had the same peak square under the same patch location and motion direction, that square was considered an initiation dendritic square.

### Topological distance between dendritic squares within the same neuron

We calculated the topological distances between dendritic squares within the same LINs using customized codes based on the A* search algorithm (https://www.mathworks.com/matlabcentral/fileexchange/26248-a-a-star-search-for-path-planning-tutorial). Dendritic square masks were first summed, dilated by 1.05 µm and manually cropped into ’backbones’. The two ends of each backbone corresponded to dendritic branching points or dendritic terminals, and no branching point was included in the middle of a backbone. All dendritic square masks were assigned to one of the backbones. The two ends of each backbone were then detected using the MATLAB built-in function ‘bwskel’. Next, A* searching was performed between all pairs of ends, and also between each dendritic square and the two ends of the backbone to which the square belonged. For pairs of dendritic squares from the same backbone, we performed A* searching to directly measure their topological distances. For pairs on different backbones, we added up the A* distances between each dendritic square and the ends of its respective backbone, as well as the A* distances between the ends of the backbones.

### QUANTIFICATION AND STATISTICAL ANALYSIS

Statistical tests were conducted using MATLAB. Non-parametric tests were used for comparing two frequency distributions (Chi-squared test of independence), and two related groups (Wilcoxon signed-rank test). Pearson correlation was used for quantifying the linear relationship between two variables. Linear mixed-effects models (’fitlme’ function in MATLAB)^84^ were used for inferential tests with nested data points. p < 0.05 was considered significant. Additional details on sample sizes, statistical test, significant levels for each experiment can be found in figure legends, Results and Methods. All acquired data that met the stated criteria were included in the analyses.

## DATA AND SOFTWARE AVAILABILITY

Requests for analyses and raw data on calcium imaging results may be made to the Lead Contact, Liang Liang.

## References

1. Yoshida, K., Watanabe, D., Ishikane, H., Tachibana, M., Pastan, I., and Nakanishi, S. (2001). A Key Role of Starburst Amacrine Cells in Originating Retinal Directional Selectivity and Optokinetic Eye Movement. Neuron 30, 771–780. 10.1016/s0896-6273(01)00316-6.

2. Fried, S.I., Münch, T.A., and Werblin, F.S. (2002). Mechanisms and circuitry underlying directional selectivity in the retina. Nature 420, 411–414. 10.1038/nature01179.

3. Demb, J.B. (2007). Cellular Mechanisms for Direction Selectivity in the Retina. Neuron 55, 179–186. 10.1016/j.neuron.2007.07.001.

4. Lee, S.-H., Kwan, A.C., Zhang, S., Phoumthipphavong, V., Flannery, J.G., Masmanidis, S.C., Taniguchi, H., Huang, Z.J., Zhang, F., Boyden, E.S., et al. (2012). Activation of specific interneurons improves V1 feature selectivity and visual perception. Nature 488, 379–383. 10.1038/nature11312.

5. Wilson, N.R., Runyan, C.A., Wang, F.L., and Sur, M. (2012). Division and subtraction by distinct cortical inhibitory networks in vivo. Nature 488, 343–348. 10.1038/nature11347.

6. Mauss, A.S., Vlasits, A., Borst, A., and Feller, M. (2016). Visual Circuits for Direction Selectivity. Annu. Rev. Neurosci. 40, 1–20. 10.1146/annurev-neuro-072116-031335.

7. Wei, W. (2018). Neural Mechanisms of Motion Processing in the Mammalian Retina. Annu Rev Vis Sc 4, 1–28. 10.1146/annurev-vision-091517-034048.

8. Pouille, F., Marin-Burgin, A., Adesnik, H., Atallah, B.V., and Scanziani, M. (2009). Input normalization by global feedforward inhibition expands cortical dynamic range. Nat. Neurosci. 12, 1577–1585. 10.1038/nn.2441.

9. Olsen, S.R., Bhandawat, V., and Wilson, R.I. (2010). Divisive Normalization in Olfactory Population Codes. Neuron 66, 287–299. 10.1016/j.neuron.2010.04.009.

10. Wang, X., Sommer, F.T., and Hirsch, J.A. (2011). Inhibitory circuits for visual processing in thalamus. Curr Opin Neurobiol 21, 726–733. 10.1016/j.conb.2011.06.004.

11. Kuffler, S.W. (1953). DISCHARGE PATTERNS AND FUNCTIONAL ORGANIZATION OF MAMMALIAN RETINA. J Neurophysiol 16, 37–68. 10.1152/jn.1953.16.1.37.

12. Hubel, D.H., and Wiesel, T.N. (1960). Receptive fields of optic nerve fibres in the spider monkey. J. Physiol. 154, 572–580. 10.1113/jphysiol.1960.sp006596.

13. Packer, O.S., Verweij, J., Li, P.H., Schnapf, J.L., and Dacey, D.M. (2010). Blue-Yellow Opponency in Primate S Cone Photoreceptors. J. Neurosci. 30, 568–572. 10.1523/jneurosci.4738-09.2010.

14. Pouille, F., and Scanziani, M. (2001). Enforcement of Temporal Fidelity in Pyramidal Cells by Somatic Feed-Forward Inhibition. Science 293, 1159–1163. 10.1126/science.1060342.

15. Wehr, M., and Zador, A.M. (2003). Balanced inhibition underlies tuning and sharpens spike timing in auditory cortex. Nature 426, 442–446. 10.1038/nature02116.

16. Blitz, D.M., and Regehr, W.G. (2005). Timing and Specificity of Feed-Forward Inhibition within the LGN. Neuron 45, 917–928. 10.1016/j.neuron.2005.01.033.

17. Zhu, Y., Qiao, W., Liu, K., Zhong, H., and Yao, H. (2015). Control of response reliability by parvalbumin-expressing interneurons in visual cortex. Nat. Commun. 6, 6802. 10.1038/ncomms7802.

18. Famiglietti, E.V. (1991). Synaptic organization of starburst amacrine cells in rabbit retina: Analysis of serial thin sections by electron microscopy and graphic reconstruction. J. Comp. Neurol. 309, 40–70. 10.1002/cne.903090105.

19. Grimes, W.N., Zhang, J., Graydon, C.W., Kachar, B., and Diamond, J.S. (2010). Retinal Parallel Processors: More than 100 Independent Microcircuits Operate within a Single Interneuron. Neuron 65, 873–885. 10.1016/j.neuron.2010.02.028.

20. Lei, W., Clark, D.A., and Demb, J.B. (2024). Compartmentalized pooling generates orientation selectivity in wide-field amacrine cells. Proc. Natl. Acad. Sci. 121, e2411130121. 10.1073/pnas.2411130121.

21. Rall, W., Shepherd, G.M., Reese, T.S., and Brightman, M.W. (1966). Dendrodendritic synaptic pathway for inhibition in the olfactory bulb. Exp. Neurol. 14, 44–56. 10.1016/0014-4886(66)90023-9.

22. Whyland, K.L., Slusarczyk, A.S., and Bickford, M.E. (2020). GABAergic cell types in the superficial layers of the mouse superior colliculus. J. Comp. Neurol. 528, 308–320. 10.1002/cne.24754.

23. Guillery, R.W. (1969). The organization of synaptic interconnections in the laminae of the dorsal lateral geniculate nucleus of the cat. Zeitschrift Für Zellforschung Und Mikroskopische Anatomie 96, 1–38. 10.1007/bf00321474.

24. Rafols, J.A., and Valverde, F. (1973). The structure of the dorsal lateral geniculate nucleus in the mouse. A golgi and electron microscopic study. J. Comp. Neurol. 150, 303–331. 10.1002/cne.901500305.

25. Hamos, J.E., Horn, S.C.V., Raczkowski, D., Uhlrich, D.J., and Sherman, S.M. (1985). Synaptic connectivity of a local circuit neurone in lateral geniculate nucleus of the cat. Nature 317, 618–621. 10.1038/317618a0.

26. Montero, V.M. (1986). Localization of γ-aminobutyric acid (GABA) in type 3 cells and demonstration of their source to f2 terminals in the cat lateral geniculate nucleus: A golgi-electron-microscopic GABA-immunocytochemical study. J. Comp. Neurol. 254, 228–245. 10.1002/cne.902540207.

27. Sherman, S.M. (2004). Interneurons and triadic circuitry of the thalamus. Trends Neurosci 27, 670–675. 10.1016/j.tins.2004.08.003.

28. Morgan, J.L., and Lichtman, J.W. (2020). An Individual Interneuron Participates in Many Kinds of Inhibition and Innervates Much of the Mouse Visual Thalamus. Neuron. 10.1016/j.neuron.2020.02.001.

29. Euler, T., Detwiler, P.B., and Denk, W. (2002). Directionally selective calcium signals in dendrites of starburst amacrine cells. Nature 418, 845–852. 10.1038/nature00931.

30. Balu, R., Pressler, R.T., and Strowbridge, B.W. (2007). Multiple Modes of Synaptic Excitation of Olfactory Bulb Granule Cells. J. Neurosci. 27, 5621–5632. 10.1523/jneurosci.4630-06.2007.

31. Zhou, Z.J., and Fain, G.L. (1996). Starburst amacrine cells change from spiking to nonspiking neurons during retinal development. Proc. Natl. Acad. Sci. 93, 8057–8062. 10.1073/pnas.93.15.8057.

32. Park, S.J., Lei, W., Pisano, J., Orpia, A., Minehart, J., Pottackal, J., Hanke-Gogokhia, C., Zapadka, T.E., Clarkson-Paredes, C., Popratiloff, A., et al. (2024). Molecular identification of wide-field amacrine cells in mouse retina that encode stimulus orientation. 10.7554/elife.94985.1.

33. Egger, V., Svoboda, K., and Mainen, Z.F. (2003). Mechanisms of Lateral Inhibition in the Olfactory Bulb: Efficiency and Modulation of Spike-Evoked Calcium Influx into Granule Cells. J. Neurosci. 23, 7551–7558. 10.1523/jneurosci.23-20-07551.2003.

34. Pressler, R.T., and Strowbridge, B.W. (2019). Functional Specialization of Interneuron Dendrites: Identification of Action Potential Initiation Zone in Axonless Olfactory Bulb Granule Cells. J. Neurosci. 39, 9674–9688. 10.1523/jneurosci.1763-19.2019.

35. Suresh, V., Çiftçioğlu, U.M., Wang, X., Lala, B.M., Ding, K.R., Smith, W.A., Sommer, F.T., and Hirsch, J.A. (2016). Synaptic Contributions to Receptive Field Structure and Response Properties in the Rodent Lateral Geniculate Nucleus of the Thalamus. J Neurosci 36, 10949–10963. 10.1523/jneurosci.1045-16.2016.

36. Gorin, A.S., Miao, Y., Ahn, S., Suresh, V., Su, Y., Ciftcioglu, U.M., Sommer, F.T., and Hirsch, J.A. (2023). Local interneurons in the murine visual thalamus have diverse receptive fields and can provide feature selective inhibition to relay cells. bioRxiv, 2023.08.10.549394. 10.1101/2023.08.10.549394.

37. Sherman, S.M., and Guillery, R.W. (1996). Functional organization of thalamocortical relays. J. Neurophysiol. 76, 1367–1395. 10.1152/jn.1996.76.3.1367.

38. Usrey, W.M., and Alitto, H.J. (2015). Visual Functions of the Thalamus. Annu Rev Vis Sc 1, 351–371. 10.1146/annurev-vision-082114-035920.

39. Liang, L., Fratzl, A., Reggiani, J.D.S., Mansour, O.E., Chen, C., and Andermann, M.L. (2020). Retinal Inputs to the Thalamus Are Selectively Gated by Arousal. Curr Biol 30, 3923–3934.e9. 10.1016/j.cub.2020.07.065.

40. Liang, L., and Chen, C. (2020). Organization, Function, and Development of the Mouse Retinogeniculate Synapse. Annu Rev Vis Sc 6, 261–285. 10.1146/annurev-vision-121219-081753.

41. Fei, Y., Luh, M.Y., Ontiri, A., Ghauri, D., Hu, W., and Liang, L. (2025). Coordination of distinct sources of excitatory inputs enhances motion selectivity in the mouse visual thalamus. Neuron. 10.1016/j.neuron.2025.07.015.

42. Fisher, T.G., Alitto, H.J., and Usrey, W.M. (2017). Retinal and Nonretinal Contributions to Extraclassical Surround Suppression in the Lateral Geniculate Nucleus. J Neurosci 37, 226–235. 10.1523/jneurosci.1577-16.2016.

43. Tschetter, W.W., Govindaiah, G., Etherington, I.M., Guido, W., and Niell, C.M. (2018). Refinement of Spatial Receptive Fields in the Developing Mouse Lateral Geniculate Nucleus Is Coordinated with Excitatory and Inhibitory Remodeling. J Neurosci 38, 4531– 4542. 10.1523/jneurosci.2857-17.2018.

44. Crandall, S.R., and Cox, C.L. (2012). Local Dendrodendritic Inhibition Regulates Fast Synaptic Transmission in Visual Thalamus. J. Neurosci. 32, 2513–2522. 10.1523/jneurosci.4402-11.2012.

45. Acuna-Goycolea, C., Brenowitz, S.D., and Regehr, W.G. (2008). Active Dendritic Conductances Dynamically Regulate GABA Release from Thalamic Interneurons. Neuron 57, 420–431. 10.1016/j.neuron.2007.12.022.

46. Cox, C.L., Zhou, Q., and Sherman, S.M. (1998). Glutamate locally activates dendritic outputs of thalamic interneurons. Nature 394, 478–482. 10.1038/28855.

47. Cox, C.L., and Sherman, S.M. (2000). Control of Dendritic Outputs of Inhibitory Interneurons in the Lateral Geniculate Nucleus. Neuron 27, 597–610. 10.1016/s0896-6273(00)00069-6.

48. Pressler, R.T., and Regehr, W.G. (2013). Metabotropic Glutamate Receptors Drive Global Persistent Inhibition in the Visual Thalamus. J. Neurosci. 33, 2494–2506. 10.1523/jneurosci.3458-12.2013.

49. Wang, X., Vaingankar, V., Sanchez, C.S., Sommer, F.T., and Hirsch, J.A. (2011). Thalamic interneurons and relay cells use complementary synaptic mechanisms for visual processing. Nat Neurosci 14, 224–231. 10.1038/nn.2707.

50. Müllner, F.E., and Roska, B. (2024). Individual thalamic inhibitory interneurons are functionally specialized toward distinct visual features. Neuron. 10.1016/j.neuron.2024.06.001.

51. Liang, L., Fratzl, A., Goldey, G., Ramesh, R.N., Sugden, A.U., Morgan, J.L., Chen, C., and Andermann, M.L. (2018). A Fine-Scale Functional Logic to Convergence from Retina to Thalamus. Cell 173, 1343–1355.e24. 10.1016/j.cell.2018.04.041.

52. Reggiani, J.D.S., Jiang, Q., Barbini, M., Lutas, A., Liang, L., Fernando, J., Deng, F., Wan, J., Li, Y., Chen, C., et al. (2023). Brainstem serotonin neurons selectively gate retinal information flow to thalamus. Neuron 111, 711–726.e11. 10.1016/j.neuron.2022.12.006.

53. Bakken, T.E., Velthoven, C.T. van, Menon, V., Hodge, R.D., Yao, Z., Nguyen, T.N., Graybuck, L.T., Horwitz, G.D., Bertagnolli, D., Goldy, J., et al. (2021). Single-cell and single-nucleus RNA-seq uncovers shared and distinct axes of variation in dorsal LGN neurons in mice, non-human primates, and humans. Elife 10, e64875. 10.7554/elife.64875.

54. Reese, B.E. (1988). ‘Hidden lamination’ in the dorsal lateral geniculate nucleus: the functional organization of this thalamic region in the rat. Brain Res Rev 13, 119–137. 10.1016/0165-0173(88)90017-3.

55. Cruz-Martín, A., El-Danaf, R.N., Osakada, F., Sriram, B., Dhande, O.S., Nguyen, P.L., Callaway, E.M., Ghosh, A., and Huberman, A.D. (2014). A dedicated circuit links direction-selective retinal ganglion cells to the primary visual cortex. Nature 507, 358–361. 10.1038/nature12989.

56. Dhande, O.S., Stafford, B.K., Lim, J.-H.A., and Huberman, A.D. (2015). Contributions of Retinal Ganglion Cells to Subcortical Visual Processing and Behaviors. Annu Rev Vis Sc 1, 291–328. 10.1146/annurev-vision-082114-035502.

57. Rodieck, R.W. (1967). Receptive Fields in the Cat Retina: A New Type. Science 157, 90–92. 10.1126/science.157.3784.90.

58. Piscopo, D.M., El-Danaf, R.N., Huberman, A.D., and Niell, C.M. (2013). Diverse Visual Features Encoded in Mouse Lateral Geniculate Nucleus. J Neurosci 33, 4642–4656. 10.1523/jneurosci.5187-12.2013.

59. Tien, N.-W., Pearson, J.T., Heller, C.R., Demas, J., and Kerschensteiner, D. (2015). Genetically Identified Suppressed-by-Contrast Retinal Ganglion Cells Reliably Signal Self-Generated Visual Stimuli. J Neurosci 35, 10815–10820. 10.1523/jneurosci.1521-15.2015.

60. Mukamel, E.A., Nimmerjahn, A., and Schnitzer, M.J. (2009). Automated Analysis of Cellular Signals from Large-Scale Calcium Imaging Data. Neuron 63, 747–760. 10.1016/j.neuron.2009.08.009.

61. Pnevmatikakis, E.A., Soudry, D., Gao, Y., Machado, T.A., Merel, J., Pfau, D., Reardon, T., Mu, Y., Lacefield, C., Yang, W., et al. (2016). Simultaneous Denoising, Deconvolution, and Demixing of Calcium Imaging Data. Neuron 89, 285–299. 10.1016/j.neuron.2015.11.037.

62. Giovannucci, A., Friedrich, J., Gunn, P., Kalfon, J., Brown, B.L., Koay, S.A., Taxidis, J., Najafi, F., Gauthier, J.L., Zhou, P., et al. (2019). CaImAn an open source tool for scalable calcium imaging data analysis. Elife 8, e38173. 10.7554/elife.38173.

63. Williams, A.H., Kim, T.H., Wang, F., Vyas, S., Ryu, S.I., Shenoy, K.V., Schnitzer, M., Kolda, T.G., and Ganguli, S. (2018). Unsupervised Discovery of Demixed, Low-Dimensional Neural Dynamics across Multiple Timescales through Tensor Component Analysis. Neuron 98, 1099–1115.e8. 10.1016/j.neuron.2018.05.015.

64. Bollmann, J.H., and Engert, F. (2009). Subcellular Topography of Visually Driven Dendritic Activity in the Vertebrate Visual System. Neuron 61, 895–905. 10.1016/j.neuron.2009.01.018.

65. Chen, T.-W., Wardill, T.J., Sun, Y., Pulver, S.R., Renninger, S.L., Baohan, A., Schreiter, E.R., Kerr, R.A., Orger, M.B., Jayaraman, V., et al. (2013). Ultrasensitive fluorescent proteins for imaging neuronal activity. Nature 499, 295–300. 10.1038/nature12354.

66. Runyan, C.A., and Sur, M. (2013). Response Selectivity Is Correlated to Dendritic Structure in Parvalbumin-Expressing Inhibitory Neurons in Visual Cortex. J. Neurosci. 33, 11724–11733. 10.1523/jneurosci.2196-12.2013.

67. Seung, H.S. (2024). Interneuron diversity and normalization specificity in a visual system. bioRxiv, 2024.04.03.587837. 10.1101/2024.04.03.587837.

68. Sillito, A.M., and Kemp, J.A. (1983). The influence of GABAergic inhibitory processes on the receptive field structure of X and Y cells in cat dorsal lateral geniculate nucleus (dLGN). Brain Res. 277, 63–77. 10.1016/0006-8993(83)90908-3.

69. Norton, T.T., Holdefer, R.N., and Godwin, D.W. (1989). Effects of bicuculline on receptive field center sensitivity of relay cells in the lateral geniculate nucleus. Brain Res. 488, 348–352. 10.1016/0006-8993(89)90728-2.

70. Norton, T.T., and Godwin, D.W. (1992). Chapter 10 Inhibitory GABAergic control of visual signals at the lateral geniculate nucleus. Prog. Brain Res. 90, 193–217. 10.1016/s0079-6123(08)63615-8.

71. Hong, Y.K., and Chen, C. (2011). Wiring and rewiring of the retinogeniculate synapse. Curr Opin Neurobiol 21, 228–237. 10.1016/j.conb.2011.02.007.

72. Leist, M., Datunashvilli, M., Kanyshkova, T., Zobeiri, M., Aissaoui, A., Cerina, M., Romanelli, M.N., Pape, H.-C., and Budde, T. (2016). Two types of interneurons in the mouse lateral geniculate nucleus are characterized by different h-current density. Sci Rep-uk 6, 24904. 10.1038/srep24904.

73. Charpak, S., Mertz, J., Beaurepaire, E., Moreaux, L., and Delaney, K. (2001). Odor-evoked calcium signals in dendrites of rat mitral cells. Proc. Natl. Acad. Sci. 98, 1230– 1234. 10.1073/pnas.98.3.1230.

74. Brainard, D.H. (1997). The Psychophysics Toolbox. Spat Vis 10, 433–436.

75. Friedrichsen, K., Ramakrishna, P., Hsiang, J.-C., Valkova, K., Kerschensteiner, D., and Morgan, J.L. (2022). Reconstructing neural circuits using multiresolution correlated light and electron microscopy. Front. Neural Circuits 16, 753496. 10.3389/fncir.2022.753496.

76. Arshadi, C., Günther, U., Eddison, M., Harrington, K.I.S., and Ferreira, T.A. (2021). SNT: a unifying toolbox for quantification of neuronal anatomy. Nat. Methods 18, 374–377. 10.1038/s41592-021-01105-7.

77. Bonin, V., Histed, M.H., Yurgenson, S., and Reid, R.C. (2011). Local diversity and fine-scale organization of receptive fields in mouse visual cortex. Journal of Neuroscience 31, 18506–18521. 10.1523/jneurosci.2974-11.2011.

78. Pachitariu, M., Stringer, C., Dipoppa, M., Schröder, S., Rossi, L.F., Dalgleish, H., Carandini, M., and Harris, K.D. (2017). Suite2p: beyond 10,000 neurons with standard two-photon microscopy. bioRxiv, 061507. 10.1101/061507.

79. Dipoppa, M., Ranson, A., Krumin, M., Pachitariu, M., Carandini, M., and Harris, K.D. (2018). Vision and Locomotion Shape the Interactions between Neuron Types in Mouse Visual Cortex. Neuron 98, 602–615.e8. 10.1016/j.neuron.2018.03.037.

80. Baden, T., Berens, P., Franke, K., Rosón, M.R., Bethge, M., and Euler, T. (2016). The functional diversity of retinal ganglion cells in the mouse. Nature 529, 345–350. 10.1038/nature16468.

81. Carandini, M., and Ferster, D. (2000). Membrane Potential and Firing Rate in Cat Primary Visual Cortex. J. Neurosci. 20, 470–484. 10.1523/jneurosci.20-01-00470.2000.

82. Zhao, X., Chen, H., Liu, X., and Cang, J. (2013). Orientation-selective Responses in the Mouse Lateral Geniculate Nucleus. J Neurosci 33, 12751–12763. 10.1523/jneurosci.0095-13.2013.

83. Rosón, M.R., Bauer, Y., Kotkat, A.H., Berens, P., Euler, T., and Busse, L. (2019). Mouse dLGN Receives Functional Input from a Diverse Population of Retinal Ganglion Cells with Limited Convergence. Neuron 102, 462–476.e8. 10.1016/j.neuron.2019.01.040.

84. Ferguson, K.A., Salameh, J., Alba, C., Selwyn, H., Barnes, C., Lohani, S., and Cardin, J.A. (2023). VIP interneurons regulate cortical size tuning and visual perception. Cell Reports 42, 113088. 10.1016/j.celrep.2023.113088.

